# Extracellular Particles Derived from Mesenchymal Stromal Cells Reduce *Pseudomonas aeruginosa* Lung Infection and Inflammation in Mice

**DOI:** 10.1101/2025.03.30.646208

**Authors:** Sharanya Sarkar, Roxanna Barnaby, Zachary Faber, Lily Taub, Carolyn Roche, Lily A. Charpentier, Thomas H. Hampton, Brad Vietje, Douglas J. Taatjes, Byoung-Kyu Cho, Young Ah Goo, Matthew J. Wargo, Daniel J. Weiss, Tracey L. Bonfield, Thomas J. Kelley, Bruce A. Stanton

## Abstract

The World Health Organization and the U.S. Centers for Disease Control and Prevention have reported that antibiotic resistant infections with *Pseudomonas aeruginosa* present a significant health risk world-wide. In the genetic disease Cystic Fibrosis (CF), chronic antibiotic resistant *Pseudomonas* lung infections and persistent inflammation remain the leading causes of mortality. While highly effective modulator therapy (HEMT) dramatically improves lung function in CF, they fail to eradicate chronic infections or eliminate the associated hyperinflammatory state. Thus, there is an urgent need for innovative therapies that can simultaneously eliminate antibiotic resistant *P. aeruginosa* lung infection and the attendant hyperinflammatory lung environment. Mesenchymal stromal cell-derived extracellular particles (MSC EPs) represent a promising solution, offering potent anti-inflammatory and antimicrobial properties while being safe and non-toxic. This study demonstrates using a CF mouse model of infection that MSC EPs reduce acute *P. aeruginosa* lung infection and inflammation. As the first investigation of MSC EPs in CF mice, this research underscores the dual effects of MSC EPs; reducing inflammation and bacterial burden. These findings mark an important advancement in antimicrobial therapy, addressing the unmet need for reducing antibiotic resistant infections and hyperinflammation in CF as well as other diseases with chronic, antibiotic resistant *P. aeruginosa* infections.

## INTRODUCTION

The World Health Organization and the U.S. Center for Disease Control and Prevention have reported that antibiotic resistant infections with *Pseudomonas aeruginosa* (*P. aeruginosa*) present a significant health risk world-wide. *P. aeruginosa* is the fifth most common cause of hospital-acquired infections, and in the U.S. alone 32,600 hospitalized patients had antibiotic resistant *P. aeruginosa* infections in 2017 (1). More than 50% of adults with the genetic disease Cystic Fibrosis (CF) are chronically infected with antibiotic resistant strains of *P. aeruginosa* (2, 3), which is the primary contributor to morbidity and mortality in people with CF (pwCF) (4, 5).

Chronic infection in CF initiates a sustained inflammatory response that fails to resolve the infection and leads to progressive lung damage (6). A significant advancement in CF research has been the development of CFTR modulators, which are small molecules designed to correct protein misfolding or enhance channel gating to facilitate apical chloride and bicarbonate secretion (2). While highly effective modulator therapy (HEMT) has shown promising results in improving lung function and reducing hospitalization rates in pwCF, it does not eradicate bacterial lung infections or mitigate the hyperinflammatory response to chronic *P. aeruginosa* infections (7, 8). Thus, there is a pressing need to develop more effective therapies to eliminate chronic antibiotic resistant *P. aeruginosa* lung infections.

Therapeutic anti-inflammatory medications have demonstrated efficacy in mitigating the decline in lung function associated with chronic *P. aeruginosa* infection, especially in pwCF. However, existing anti-inflammatory drugs elicit liver, pancreas, and kidney complications rendering them inadequate as therapeutics for sustained inflammation associated with chronic infection (9). Consequently, there is also a need for novel treatments to address inflammation in people with chronic *P. aeruginosa* infections. Given the substantial global burden of bacterial antimicrobial resistance (estimated to reach US$1 trillion/year by 2050), developing new antibacterial and anti-inflammatory therapies holds promise for addressing various respiratory infections beyond pwCF (10).

Mesenchymal stromal cells (MSCs), precursor cells originating from bone marrow, adipose tissue, or cord blood, possess immunomodulatory and antibacterial characteristics and have been investigated for various acute and chronic inflammatory lung conditions (11–14). MSCs exhibit low immunogenicity, allowing for their utilization as an allogeneic cell source without the risk of rejection (15). Additionally, MSCs demonstrate favorable anti-inflammatory and tissue regenerative properties (16–18). MSCs themselves and the soluble factors they secrete impede the growth of *P. aeruginosa*, *Staphylococcus aureus*, and *Streptococcus pneumoniae* due to the secretion of antimicrobial peptides (19). These desirable attributes have rendered MSCs a subject of great interest to researchers, with ongoing investigations spanning over 1,200 clinical trials (11). The established safety and lack of toxicity of MSCs have led to regulatory approvals for MSC-based therapies in several countries (20). Efforts in the CF setting have demonstrated the safety of allogeneic human MSCs in a phase 1 study involving 15 adult pwCF (21).

Researchers initially believed that MSCs exerted their effects solely through direct cellular interactions (22) until two seminal papers in the field unveiled that extracellular vesicles (EVs) released by MSCs also possess the properties of their parent cells (23, 24). MSC-derived EVs contain an array of biomolecules, including proteins, mRNA, lipids, DNAs, and miRNAs (25), and have demonstrated paracrine effects in various animal models of injury as extensively reviewed elsewhere (22, 26–28). Additionally, MSC EVs have shown promise in mitigating inflammation and tissue damage associated with pulmonary diseases as comprehensively discussed in recent reviews (25, 29). Beyond their anti-inflammatory roles, MSC EVs exhibit antimicrobial properties (30, 31). MSC EVs offer several advantages over MSCs themselves. First, EVs are less prone to inflammatory environmental alterations than MSCs (32). Second, their nano-scale size allows them to traverse mucus layers and penetrate antibiotic resistant bacterial biofilms effectively (33, 34). Third, their smaller size and reduced expression of MHC molecules make them less susceptible to immune rejection compared to their parent cells (35). The multitude of these benefits, coupled with their proven safety in clinical trials, position MSC EVs as promising candidates for therapeutic intervention (36, 37). We have shown previously that the miRNA let-7b-5p in EVs secreted by primary human bronchial epithelial cells (HBECs), when combined with sub-minimum inhibitory concentrations (MIC) of aztreonam or carbenicillin, antibiotics commonly used in people with CF (pwCF) effectively eradicated the formation of antibiotic-resistant *P. aeruginosa* biofilms *in vitro* (38). This combination also increased the antibiotic susceptibility of both mucoid and non-mucoid clinical strains of *P. aeruginosa* (38).

Because MSC EVs contain negligible levels of let-7b-5p (39, 40), and let-7b-5p has antibacterial (38, 41, 42) and anti-inflammatory properties (43–45), the goal of this study was to engineer MSC EVs to contain let-7b-5p as a therapeutic approach to reduce infection and inflammation using CF mice as a model of infection (39, 46). To our knowledge there are no studies or registered clinical trials involving let-7 family members to treat antibiotic resistant *P. aeruginosa* lung infections (47), highlighting a significant knowledge gap. To align with MISEV guidelines, we refer to our MSC-derived extracellular vesicles (EVs) as MSC-derived extracellular particles (EPs) throughout the manuscript.

Our data demonstrates that MSC EPs, even without antibiotics, decrease the burden of a clinical mucoid isolate of *P. aeruginosa* and immune cells and modulate cytokine levels in the lungs of CF mice, thereby highlighting their potential as a novel anti-infective and anti-inflammatory candidate for antibiotic resistant *P. aeruginosa* lung infections.

## MATERIALS AND METHODS

### Isolation of MSC-derived extracellular particles (MSC EPs) and miRNA transfection of MSCs

Human Bone Marrow-Derived MSCs were obtained from the ATCC (Cat # PCS-500-012). MSCs were cultured according to the manufacturer’s instructions using the ATCC media (2% FBS: up to passage 4). MSCs were transfected with a miRNA mimic negative control (NC, that has no known targets in mice or *P. aeruginosa*) or miRNA let-7b-5p, obtained from Ambion (Austin TX) using Lipofectamine RNAiMAX Transfection Reagent (Invitrogen, Waltham, MA). To determine if lipofectamine had effects on the results, we also included a lipofectamine-only control (without miRNA) in our EV isolation workflow. Two days before transfection, MSC were seeded at 1×10^6^ per T-75 flask. Once ∼80% confluent, the media was switched to supplement-free (FBS-Free) media, and cells were transfected with a 20 μM stock of let-7b-5p (Ambion catalog #4464066) or the negative control miRNA (Ambion catalog #4464058). A mixture of RNAiMax (1:1000 vol/vol of reagent to total cell media), OptiMem (Invitrogen catalog #31985070), and miRNA was allowed to sit for 5 minutes and then added dropwise to the flasks containing MSCs. One flask of MSCs was not transfected at all (referred to as control or untreated MSCs). The media were harvested 24 hours after the transfection. EPs were purified from the MSC culture supernatants using the ExoQuick-TC EV isolation kit (#EXOTC50A-1, Systems Biosciences Palo Alto, CA). Briefly, suspended cells were removed from MSC culture supernatants by centrifugation at 3000 g for 15 min, and 12 mL of supernatants were concentrated to ∼200 – 400 µL with a 30 kDa Amicon Ultra Centrifugal Filter (Millipore, Billerica, MA). The concentrate was mixed with 10 μL of ExoQuick-TC polymer per 50 µL concentrated media. Following overnight incubation at 4°C, EPs were isolated by centrifugation (1,500 g, 30 min) and resuspended in a final volume of 150 μL PBS. MSC growth media not conditioned by cells was subjected to the same isolation process and was termed process control (PC) to account for any effect of the cell culture media and possible contamination that may be introduced during the EP isolation process.

### qRT-PCR to assess let-7b-5p loading in MSC EPs

RNA was isolated from EPs (10 μL total EP volume) using the miRNeasy Mini Kit (Qiagen, catalog #1038703) according to the manufacturer’s instructions and eluted in RNase-free water. cDNA was made using a Taqman miRNA Kit (Invitrogen, catalog #4366597). Let7b-5p specific primers were obtained from Invitrogen (catalog #4395446). In preliminary studies U6 was selected as an appropriate reference gene for qRT-PCR, since U6 (Invitrogen, catalog #4398988) was not affected by experimental treatments. To assess whether let-7b-5p was internal to the EPs, the EPs were incubated with 50ug/mL Proteinase K (Thermo Scientific, catalog # EO0491) with or without 1% Triton-X 100 (Bio-Rad, catalog #161-0407) at 65°C for 5 minutes, followed by 80°C for 15 minutes to deactivate the Proteinase K. EPs were briefly cooled on ice before adding RNase A/T1 Mix (Thermo Scientific, catalog #EN0551) at 5 U/mL RNase A and 200 U/mL RNase T1 and incubated at 37°C for 60 minutes. Samples were immediately submitted to the above workflow for RNA isolation and qRT-PCR.

### Characterization of MSC EPs

MSC EPs were characterized via Exosome Analytical Services provided by RoosterBio Inc.; (Frederick, MD) as described and reported in our previous work (48).

### Proteomics sample preparation

Proteins were isolated from EPs secreted by human MSCs transfected with a negative control siRNA (Invitrogen), lipofectamine alone, and let-7b-5p, along with an untreated control. For protein isolation, lysis buffer (0.5% SDS, 50mM AmBic, 50mM NaCl, HALT Protease Inhibitor) was added to MSC EP samples, followed by sonication. Samples were purified by trichloroacetic acid precipitation. After pelleting and washing with ice-cold acetone, the resulting protein pellet was resuspended in 8 M urea and 0.4 M ammonium bicarbonate, reduced with 4 mM dithiothreitol, and alkylated with 18 mM iodoacetamide. The solution was then diluted to <2 M urea, and 1 µg of trypsin was added for overnight digestion at 37°C. The resulting peptides were desalted using C18 solid-phase extraction spin columns, and eluates were dried under vacuum using a SpeedVac concentrator.

### Mass spectrometry analysis

Peptides were analyzed by LC–MS/MS using a Vanquish Neo UHPLC system coupled to an Exploris 240 Orbitrap mass spectrometer (Thermo Fisher Scientific). Samples were loaded onto a Neo trap cartridge coupled with an analytical C18 column (75 µm × 25 mm, IonOpticks) and separated using a 60-minute linear gradient of solvent A (0.1% formic acid in water) and solvent B (0.1% formic acid in acetonitrile). Full MS scans were acquired at a resolution of 60,000 with an AGC target of 300% and a maximum injection time of 25 ms. For DIA acquisition, the precursor range was set to 380–980 m/z with a 10 m/z isolation window. MS2 fragmentation was performed using 28% HCD collision energy in the Orbitrap at a resolution of 15,000, with an AGC target of 2000% and a maximum injection time of 40 ms. Data processing was performed using Spectronaut 18 (Biognosys AG). Raw data files were analyzed using directDIA™ against the human SwissProt database. Search parameters included trypsin digestion with cleavage after K or R, allowance for up to two missed cleavages, carbamidomethylation of cysteine (static modification), and variable modifications including oxidized methionine and Protein N-terminal acetylation. Peptide spectrum matches (PSMs), peptides, and protein groups were identified using a false discovery rate (FDR) threshold of 0.01. Data was plotted using R v4.4.0 and tidyverse v2.0.0.

### miRNA-seq analysis of MSC EPs

Studies were also conducted to examine the miRNA content of MSC EPs by smRNA-seq, with the goal to determine the effect of transfection on the miRNA content of MSC EPs and to identify other miRNAs in MSC EPs that may have gene targets in *P. aeruginosa.* RNA was isolated from 10 µL of either control MSC EPs (no transfection), negative control miRNA transfected MSC EPs (NC MSC EPs) or let-7b-5p transfected MSC EPs (let-7b-5p MSC EPs). In some cases, RNA concentrations were too low to quantify, thus, the entire volume of extracted RNA was desiccated to 5 µL and used as input into the Qiaseq smRNA library preparation kit (Qiagen). For all samples, the workflow was performed according to manufacturer’s instructions, using 24 cycles for final library PCR. Blinded samples were uniquely indexed and pooled for sequencing on a Nextseq200 instrument (Illumina), targeting 10 million reads per sample.

miRNA-Seq reads were first trimmed using CutAdapt v4.0 with trimming parameters “-m 1, --nextseq-trim=30, -q 30, --max-n 0.8”. Unique molecular identifiers (UMIs) were then extracted from trimmed reads and appended to read named using UMI-tools extract (v1.1.5). Reads were aligned to human miRNA sequences from miRbase using Bowtie2 v2.4.4 with mapping parameters “-D 20, -R 3, -N 1, -L 12, -i, S,1,0.50”. miRbase sequences were padded by 5 base pairs at both ends with native hg38 (GRCh38.97) sequences to maximize alignment rates. Mapped reads were filtered with Samtools v1.15.1 such that matches were required to be between sixteen and twenty-eight base pairs in length, with no gaps, less than one mismatches, and a MAPQ score ≥ 20. Remaining reads were deduplicated based on UMI sequences with UMI-tools dedup (--method unique). Reads aligning to miRbase were counted using Samtools-idxstats to estimate expression levels. To facilitate detection of RNAs absent in miRbase, unmapped reads were aligned to the hg38 genome using Bowtie2 in very-sensitive-local mode, de-dupulicated, and filtered following the same criteria above, and assigned to genes using featureCounts (Subread v2.0.1). Sequencing and alignment quality control metrics were assessed using Samtools v1.15.1, FastQC v0.11.9, and Multiqc v1.12.

miRNA log_2_ fold changes (log_2_FC) between all pairwise combinations of MSC EP samples (control MSC EPs, negative control transfected MSC EPs, and let-7b-5p MSC EPs) were calculated from TMM normalized counts per million (CPM), generated from the miRbase aligned miRNA raw counts using the edgeR package (version 4.2.2) in R version 4.4.0. We used the Rocket-miR program (41) to identify pathways in *P. aeruginosa* and other ESKAPE pathogens that are potentially targeted by miRNAs of interest. Rocket-miR uses the IntaRNA program to predict interactions between human miRNAs and bacterial proteins (41, 49, 50).

IntaRNA was also used to predict human and mouse gene targets (with human genome assembly GRCh38.p14 and mouse genome assembly GRCm39) for miRNAs that had absolute log_2_FC values greater than 1 in both the let-7b-5p MSC EP and the NC MSC EP compared to control MSC EP. IntaRNA was run in IntaRNAsTar mode to optimize for whole genome searches. The top 10 gene targets by predicted interaction energy score were analyzed for functional relevance with DAVID (51, 52). We specifically focused on pathways and processes related to inflammation in both mice and humans.

### Electron microscopy of MSC EPs

EPs were visualized with negative staining transmission electron microscopy. We used EPs from untransfected MSCs, NC transfected MSCs and let-7b-5p transfected MSCs to determine if transfection alters EV morphology. A 10-microliter aliquot of each sample was applied onto a freshly glow-discharged 200-mesh nickel grid and allowed to sit for 2 minutes. After this incubation, excess fluid was gently removed with filter paper, and the grids were sequentially rinsed on seven drops of Millipore-filtered distilled water. While the grids remained slightly damp, they were placed onto a drop of Nano-W tungsten stain (Ted Pella; Product No. 09S432, Lot No. 14702). Excess stain was immediately absorbed with filter paper, and the staining step was repeated on a second drop of Nano-W, which was allowed to sit for 1 minute before being wicked away and left to dry. Imaging of the stained grids was conducted at 80 kV on a JEOL 1400 transmission electron microscope (JEOL Inc., Danvers, MA), and TIFF-format images were captured using an AMT XR11 digital camera (AMT, Woburn, MA). For diameter measurements, the electron microscopy images were processed in MetaMorph Offline image analysis software (v.7.8.0.0; Molecular Devices LLC, San Jose, CA), where 28-51 EP diameters were measured by drawing a line across each vesicle. The average diameter was subsequently calculated. The imaging and measurements of the EP diameters were conducted by an investigator blinded to the sample identities.

### Preparation of agarose-containing beads for mouse experiments

Agarose-containing beads for embedding *Pseudomonas aeruginosa* were prepared as previously described (42). In summary, five colonies of a clinical, mucoid isolate of *P. aeruginosa* tagged with mCherry (mCH PA M57-15), grown overnight on gentamicin/LB agar, were inoculated into 25 mL of LB broth and incubated at 37°C with shaking at 1 g for 20 hours. The culture was then diluted tenfold to achieve an optical density (OD) at 600 nm of approximately 0.3. 23 mL of the mCH PA M57-15/agarose/PBS mixture was added to 250 mL of sterile mineral oil in a beaker and stirred on medium-high speed. After 10 minutes, the bead/oil mixture was transferred to 15 mL of prewarmed 0.5% sodium deoxycholate/PBS and centrifuged to separate the beads from the oil. After a second wash, the beads were resuspended in 20 mL of PBS, washed four times, and imaged under the red fluorescence channel. The bead/PBS mixture was then diluted to achieve a concentration of 25,000 CFU in 50 µL, as determined by plating on gentamicin/LB agar and colony-forming unit (CFU) enumeration.

### Inoculation of Pseudomonas aeruginosa in Mice

All experiments were conducted using gut-corrected F508del/F508del Cftr^tm1Kth^ TG(FABPCFTR)1Jaw/CWR mice (C57BL/6 genetic background) that expressed CFTR in gut cells under the FABP promoter to minimize gut obstruction and enhance breeding efficiency (42). Institutional Animal Care and Use Committee (IACUC) approval from Case Western Reserve University was obtained for all protocols. Male and female mice aged 10–12 weeks were inoculated with agar beads loaded with 10^5^ CFU of the mCH PA M57-15 strain and 10^8^ MSC EPs or process control. Experiments using naïve MSC EPs, and EPs secreted by lipofectamine treated-MSC were conducted to determine lipofectamine alone might alter the EPs. Human-derived EPs have previously been evaluated in mouse models for cytotoxicity and immunogenicity, demonstrating safety (53) and efficacy across various experimental models (54, 55). The inoculum was administered to the right lung using a micro-endoscopy system (Polydiagnost, Pfaffenhofen, Germany) because preliminary experiments indicated this method increased inoculation efficiency and reduced variability. Mice were placed on warming pads post-anesthesia with isoflurane and monitored until fully recovered. Daily assessments of the inoculated mice were conducted up to the day of sample collection, following established protocols (56). Three days post-infection, the mice were euthanized via CO_2_ inhalation. After disinfecting with 70% ethanol, a midline incision was made to expose the thoracic cavity, and the diaphragm was incised to access the lungs. Bronchoalveolar lavage fluid (BALF) was collected, and both lobes of the lung were individually homogenized and cultured to assess *P. aeruginosa* growth (reported in CFU/mL). Total bacterial burden was calculated by summing colony forming units (CFU) counts from BALF, left lung, and right lung homogenates. Total white blood cells in BALF were quantified using a hemocytometer, and differential cell counts were determined through Giemsa/Miller staining. Serum and BALF cytokines were analyzed on all mice at euthanasia (day 3), using a Luminex 32-plex multianalyte cytokine array (Millipore Mouse 32-Plex Kit, EMD Millipore Corporation, Billerica, MA). All experimental protocols will be deposited in EV-TRACK to enhance transparency and interpretation of EP experiments and to increase experimental reproducibility.

### Statistical Analysis

To minimize potential bias, all analytic assays, mouse experiments and data analyses were performed by personnel blinded to the treatment. Statistical analysis was performed using GraphPad Prism (v10.3.1, San Diego, CA) and R (v 4.4.0) (57). Some mice with zero CFU counts were excluded from the analysis due to liquid handling errors. For count data such as CFUs and immune cells, negative binomial generalized Linear Models (glm.nb) were employed to compare control and EP-exposed groups. This approach was chosen because it is a robust and less biased method that appropriately accounts for the characteristics of bacterial count data, which are strictly non-negative and tend to be skewed toward larger values (58). Immune cell counts were analyzed using batch as a fixed effect, while batch was treated as a random effect for CFU counts to facilitate model convergence. For cytokine levels representing continuous data rather than counts, linear models were applied with batch included as a fixed effect. Statistical analyses were conducted using the *MASS* and *lme4* packages in R, with P-values derived accordingly. This methodological framework ensured accurate and reliable comparisons across experimental groups while accounting for batch variability.

The graphical abstract was created using the Premium version of BioRender with the following publication license: Sarkar, S. (2025) https://BioRender.com/skbgsv9

## RESULTS

### Isolation and characterization of MSC EPs

In alignment with the International Society of Extracellular Vesicles’ (ISEV) objective to enhance the robustness of EP research (59), we conducted a comprehensive analysis (particle concentration, particle size, total protein, total lipid, total miRNA, % purity) of bone marrow MSC EPs (MSCs purchased from ATCC^®^, see methods), as reported in our previous study (48). We also conducted proteomic analysis of all EPs and found the presence of EV markers (CD81, Alix) and the absence of non-EV markers (calnexin, AGO2) **(Figure 1A)** in our EV preps. We analyzed EPs derived from untreated (naïve) MSCs, lipofectamine-treated MSCs, MSCs transfected with negative control miRNA, and MSCs transfected with let-7b-5p, and observed no differences in marker profiles, indicating that lipofectamine treatment or transfection does not alter EV marker expression. **(Figure 1A)**. Since naïve MSC EPs do not contain detectable levels of let-7b-5p (39, 40), we harvested EPs from control (untreated) MSCs and MSCs transfected with either a negative control (NC) miRNA (a miRNA with no known targets in humans, mice, or *Pseudomonas*) or let-7b-5p. To ensure successful let-7b-5p loading into MSC EPs, qRT-PCR was performed on EPs using let-7b-5p specific primers, with U6 as a reference gene. The results demonstrated that while U6 levels stayed consistent among the three groups (ΔCT = 0.3), the CT (cycle threshold) value for let-7b-5p in EPs isolated from let-7b-5p transfected MSCs was lower than that from control or NC MSCs (ΔCT = 8.65). **(Figure 1B)**. This analysis demonstrated that our transfection protocol successfully loaded let-7b-5p into MSC EPs. Finally, we used a combination of treatments and qRT-PCR to determine if let-7b-5p was internalized within EPs and/or in the corona bound to AGO2 **(Figure 1C)**. RNase alone resulted in a modest increase in Ct compared to PBS (ΔCt = 1.972), indicating that some let-7b-5p miRNA is present in the corona of MSC EPs. RNase + Proteinase K treatment had no effect on let-7b-5p (ΔCt = 0.147), suggesting minimal protection of let-7b-5p by protein complexes in the corona of the EPs. In contrast, EP exposure to Triton (to enable the RNase to enter MSC EPs) led to a marked increase in Ct (ΔCt = 3.438), which was further amplified by exposure to RNase, Proteinase K and Triton (ΔCt = 9.033), demonstrating that the majority of let-7b-5p is encapsulated within EPs and that a portion of the internal miRNA is protein-associated. Moreover, the absence of AGO2 **(Figure 1A)** demonstrates that let-7b-5p in not bound to AGO2 in EPs.

**Figure 1:**
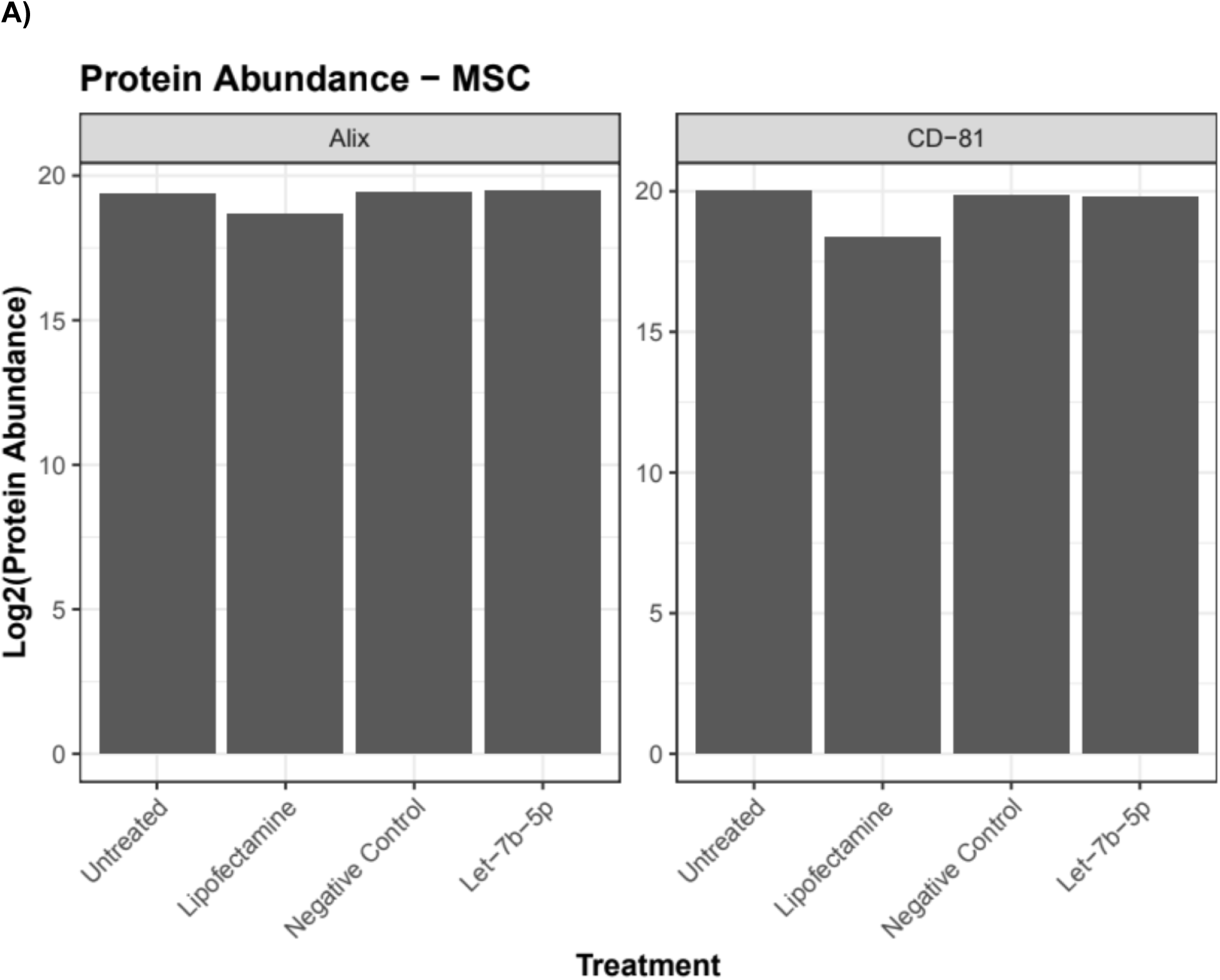
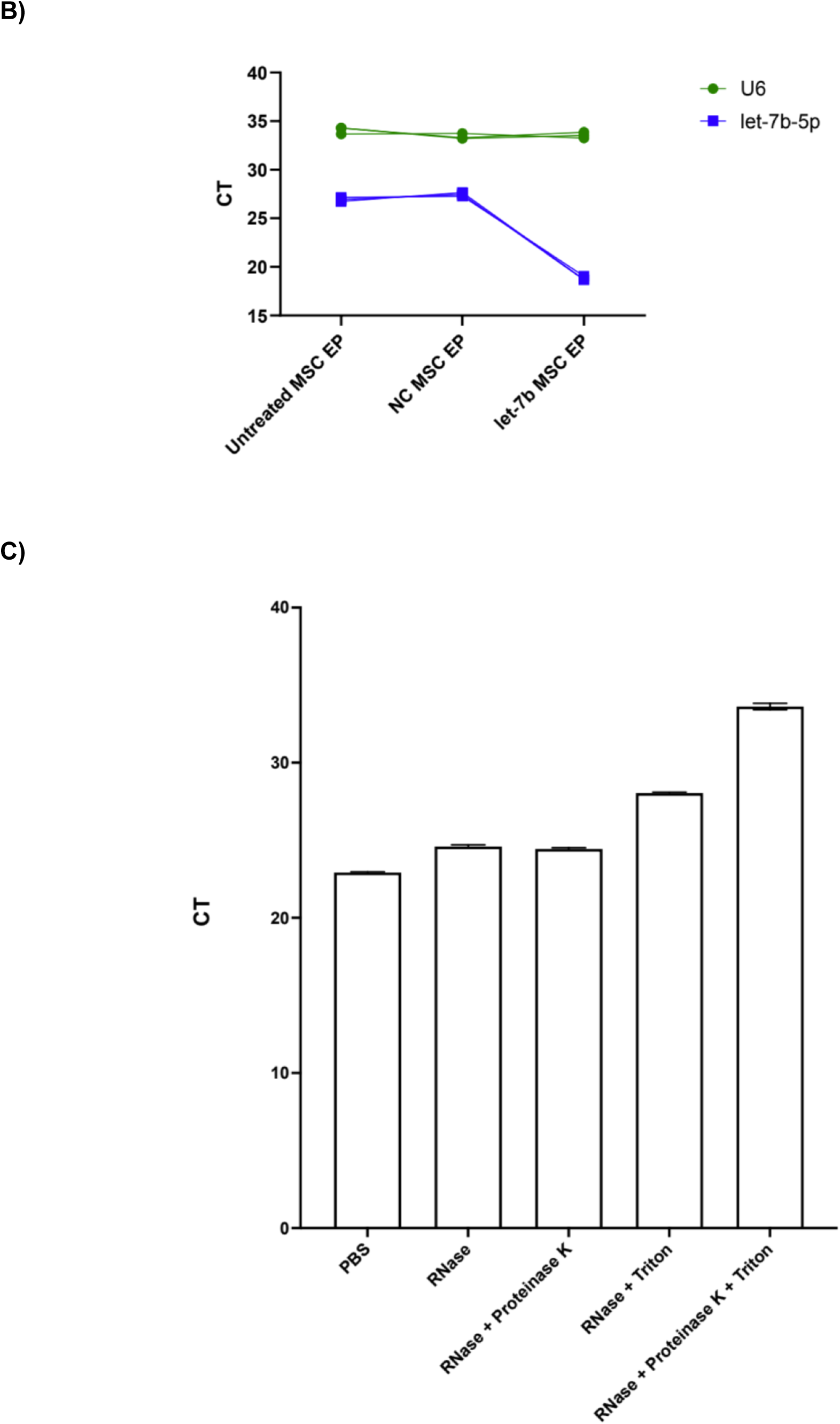
**A)** Protein marker characterization of MSC EPs via proteomics analysis. EV markers such as CD81 and ALIX were present, while non-EV markers such as calnexin and AGO2 were absent. Lipofectamine or the transfection process did not change the marker profile of EVs. **B)** qRT-PCR assays revealed a decrease in the ΔCT of let-7b-5p in EPs derived from let-7b-5p transfected MSCs compared to those from control MSCs or MSCs transfected with NC miRNA indicating successful let-7b-5p loading via parent cell transfection (ΔCT = 8.65). The reference gene, U6, was not affected by transfection (ΔCT = 0.3) among the three groups. n = 3 EP preparations; mean ± SEM depicted; NC = negative control. The colored lines indicated three experiments. **C)** RNase protection assay of let-7b-5p in MSC EPs. EP preparations were treated with PBS (control), RNase, RNase + Proteinase K, RNase + Triton, or RNase + Proteinase K + Triton prior to qPCR for let-7b-5p. A slight increase in Ct upon RNase treatment indicates a small external fraction of let-7b-5p. No change in Ct with RNase + Proteinase K suggests minimal protection of let-7b-5p by protein complexes. A marked increase in Ct following RNase + Triton reflects degradation of intravesicular let-7b-5p, with a further increase upon addition of Proteinase K indicating that a portion of the internal let-7b-5p is protein-associated. Data are presented as mean ± SEM; n = 3.

**Figure 2:**
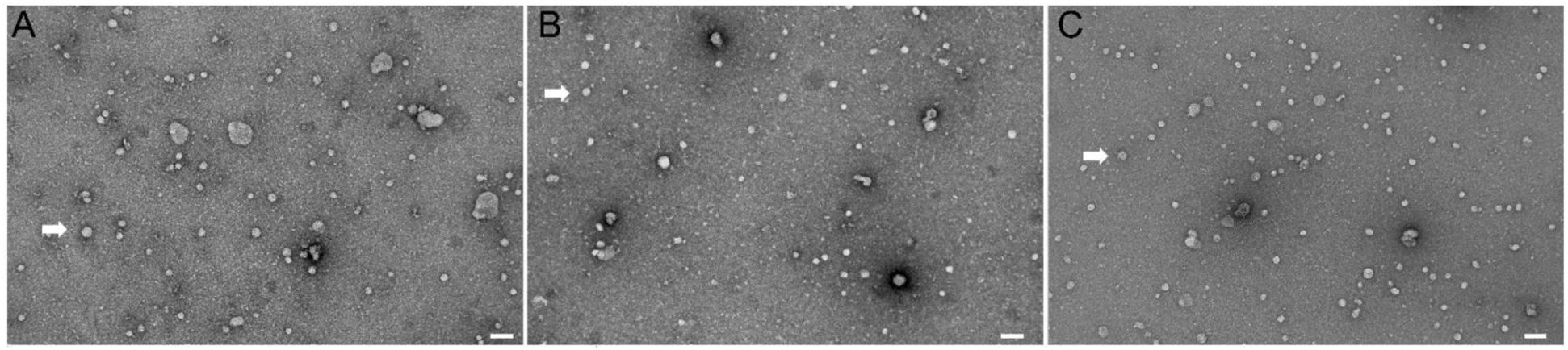
Transmission electron micrographs of A) EPs secreted by untreated MSC cells (40.8 nm ± 1.9 nm in diameter, n=35). B) EPs secreted by MSC cells treated with lipofectamine + negative control miRNA (36.5 nm ± 2.1 nm in diameter, n=28). C) EPs secreted by MSC cells treated with lipofectamine + let-7b-5p (39.5 nm ± 1.9 nm in diameter, n=51). Previously, we demonstrated that EPs were secreted by MSC, but not present in process control (48). Images A-C, scale bar 100 nm.

### Visualizing MSC EPs with Electron Microscopy

We employed high-resolution negative staining transmission electron microscopy to visualize the EPs from untransfected MSCs, NC MSCs, and let-7b-5p MSCs. Images of the EP samples identified the presence of particles and also revealed that the transfection process did not alter the morphology of EPs. The EPs had an average diameter between 37 – 40 nm, which aligns with the reported size range for EPs (42, 60, 61). Notably, there was a clear difference in the diameter of particles measured by Nanoparticle Tracking Analysis (151.1 nm) and negative staining electron microscopy (37 - 40 nm). This is not surprising given previous examples in the literature of differences in EP size measured by NTA and electron microscopy (62, 63). NTA measures light scattering and provides hydrodynamic radii, thereby overestimating EP diameter (64). The fixation process for electron microscopy can shrink particles and reduce their diameter (65).

### MSC EPs reduce the total *Pseudomonas* burden in CF mice

Before evaluating the effects of MSC EPs on *P. aeruginosa* infection and the immune response, we wanted to determine if lipofectamine altered the functional outcome of MSC EPs. For that, we ran a small pilot experiment with CF mice infected with *P. aeruginosa* treated with either naïve MSC EPs, or lipofectamine-treated MSC EPs. Mice were infected with *P. aeruginosa*-impregnated agarose beads along with either Process Control (PC), MSC EPs loaded with negative control miRNA (NC MSC EP) or MSC EPs loaded with let-7b-5p (let-7b MSC EP). After 3 days, *P. aeruginosa* CFU/mL from the harvested lung tissue and BALF were measured to assess the total bacterial burden. Both NC MSC EPs and let-7b-5p MSC EPs significantly reduced *P. aeruginosa* CFUs/mL compared to PC (p-value = 2 x 10^-16^ and 1.63 x 10^-11^, respectively) **(Figure 3B)**. Notably, the reduction in CFUs was similar in mice exposed to NC MSC EPs and let-7b-5p MSC EPs **(Figure 3B)**. Thus, the lowest concentration of lipofectamine recommended by the supplier did not alter MSC EPs’ ability to reduce total CFUs **(Figure 3A)**.

**Figure 3:**
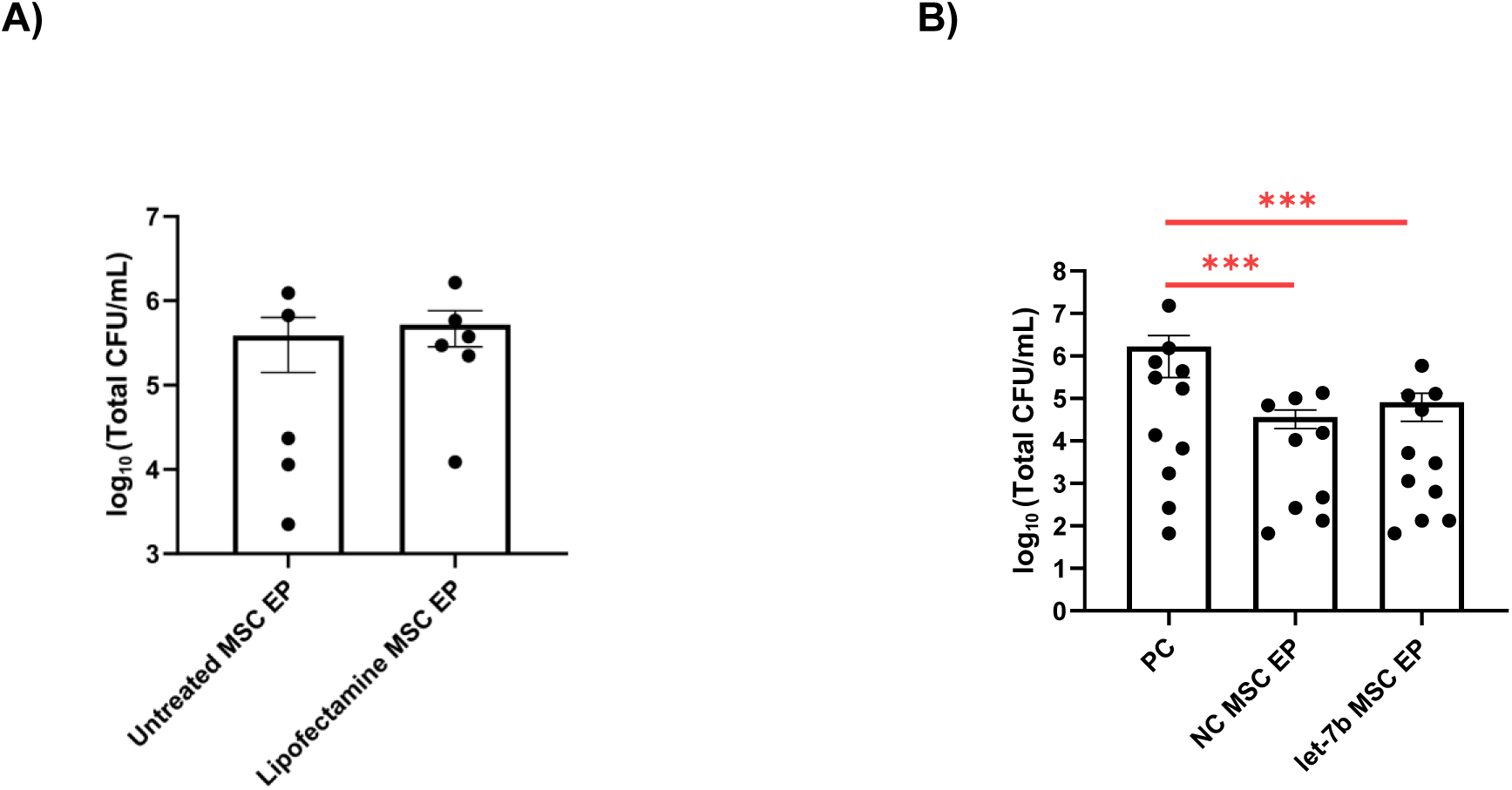
**A)** Lipofectamine exposure of MSC had no effect on total lung *Pseudomonas aeruginosa* counts (CFU/mL) in mice exposed to EPs. Compare untreated MSC EP to lipofectamine EPs (∼0.5 log_10_), indicating that lipofectamine treatment did not alter CFU counts of *P. aeruginosa*. **B)** MSC EPs loaded with negative control (NC) miRNA or let-7b-5p reduced *Pseudomonas aeruginosa* colony forming units/mL in CF mice by 2.5 log_10_ as compared to the process control (PC) group. Negative binomial generalized linear models were used to account for potential batch effects and to test the statistical significance between the PC and EP groups. Batch was modeled as a random effect to enable model convergence. Each data point represents one CF mouse. \*\*\**P* = 2 x 10^-16^ and 1.63 x 10^-11^, respectively for NC (vs PC) and let-7b (vs PC); *n* = 5 - 11 mice/group; mean ± SEM depicted. Data were compiled from 3 independent experiments (with the exception of the experiment testing effects of lipofectamine treatment, Figure 3A). PC = process control (unconditioned medium passed through the EP isolation process).

### MSC EPs modulate two key cytokines in the BALF and serum of CF mice

The effect of MSC EPs on inflammation was assessed by measuring the levels of cytokines in BALF and serum using Luminex multi-plex technology. NC MSC EPs significantly increased IL-10 in BALF (p = 0.04), compared to PC. Although the let-7b-5p MSC EPs tended to increase IL-10 compared to PC, the change was not significant (p = 0.14) **(Figure 4A)**. In serum, NC MSC EPs significantly reduced the levels of the pro-inflammatory cytokine RANTES compared to PC (p = 0.046), while the let-7b-5p MSC EP group showed a tendency to decrease RANTES but the change was not significant (p = 0.18) **(Figure 4B)**. IL-10 is an anti-inflammatory cytokine that can limit tissue damage to the host (66), a process that will be of utmost importance to people chronically infected with *P. aeruginosa.* The reduction in RANTES will reduce migration of lymphocytes and monocytes to the lungs, which would be beneficial to pwCF since recruitment of CF lymphocytes and monocytes in CF is excessive and prolonged, thereby leading to tissue damage (67). The differential change of cytokines in BALF versus serum is likely due to different levels of cytokine-binding proteins, and soluble receptors for cytokines in the two different biological fluids (68). These results demonstrate that MSC EPs, like MSCs, are anti-inflammatory and pro-resolving. Taken together, the data suggest that MSC EPs modulate cytokines in a way that would be beneficial to pwCF.

**Figure 4:**
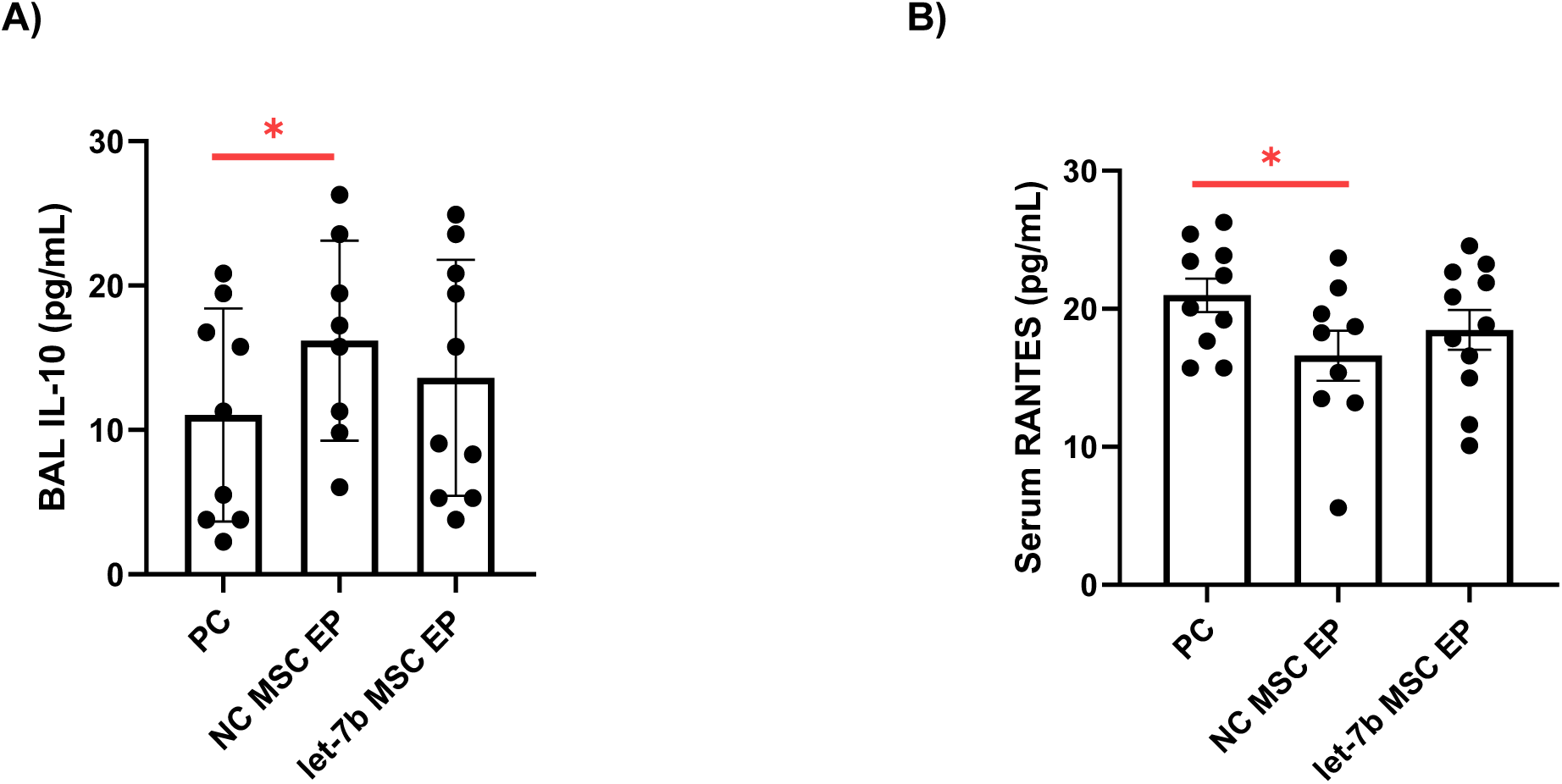
MSC EPs engineered to contain negative control (NC) miRNA significantly increased **A)** BALF IL-10 (by 46.4%) (p = 0.04) and decreased **B)** Serum RANTES (by 20.8%) as compared to the process control (PC) group (p-value = 0.046). let-7b-5p loaded MSC EPs tended to increase IL-10 in BALF (by 23.2%) (p-value = 0.14) and decrease serum RANTES (by 11.9%) (p-value = 0.18) with respect to PC mice. Linear models with batch as a fixed effect were used to account for potential batch effects and test the statistical significance between the control and EP groups. Each data point represents one CF mouse. Mice which had IL-10 and RANTES levels beyond the detection level of the multiplex platform (0.1 -1 pg/mL) were excluded from the analyses. \**P* < 0.05; *n* = 8 - 11 mice/group; mean ± SEM depicted. Data were compiled from 3 independent experiments. A power analysis of the data (at p = 0.05 and power = 0.80) revealed that an additional 140 mice are needed to obtain statistically significant changes in IL-10 and an additional 48 mice are needed to obtain significant changes in RANTES in let-7b-5p MSC EP compared to PC, making further mouse experiments impractical.

### MSC EPs reduce immune cell numbers in the BALF of CF mice

Studies were also conducted to explore the effects of MSC EPs on immune cell counts in the BALF of infected CF mice. Compared to PC, NC MSC EPs significantly reduced total cells (a summation of monocytes, lymphocytes, and neutrophils) (p = 0.00713) in BALF and let-7b-5p MSC EPs demonstrated a very strong tendency (p-value = 0.06) to reduce total cell counts **(Figure 5A)**. Both NC MSC EPs and let-7b-5p MSC EPs significantly reduced lymphocytes (p = 0.011 and 0.008 for NC (vs PC) and let-7b-5p (vs PC), respectively) **(Figure 5B)** and monocytes (p = 0.03 and 0.02 for NC (vs PC) and let-7b-5p (vs PC), respectively) **(Figure 5C)** in BALF. Neutrophil counts decreased significantly with NC MSC EPs (p = 0.018) and tended to decrease with let-7b-5p MSC EPs **(Figure 5D)**. Although not statistically significant (p-value = 0.13) the decreases in neutrophil counts was 39.9%, a decrease likely to have a positive effect since neutrophils in CF have a defect in the ability to kill *P. aeruginosa* and CF neutrophils release proteases and DNA that cause lung damage, increase mucus viscosity and thereby the ability to clear bacteria (69). The lack of significance in total cell reduction for let-7b-5p MSC EPs is attributable to the limited effect on neutrophil counts. Notably, MSC EPs in this study outperformed Human Bronchial Epithelial Cell EVs (HBEC EVs) reported in our previous study (42), as HBEC EVs failed to reduce total immune cell numbers following *P. aeruginosa* infection in CF mice.

**Figure 5:**
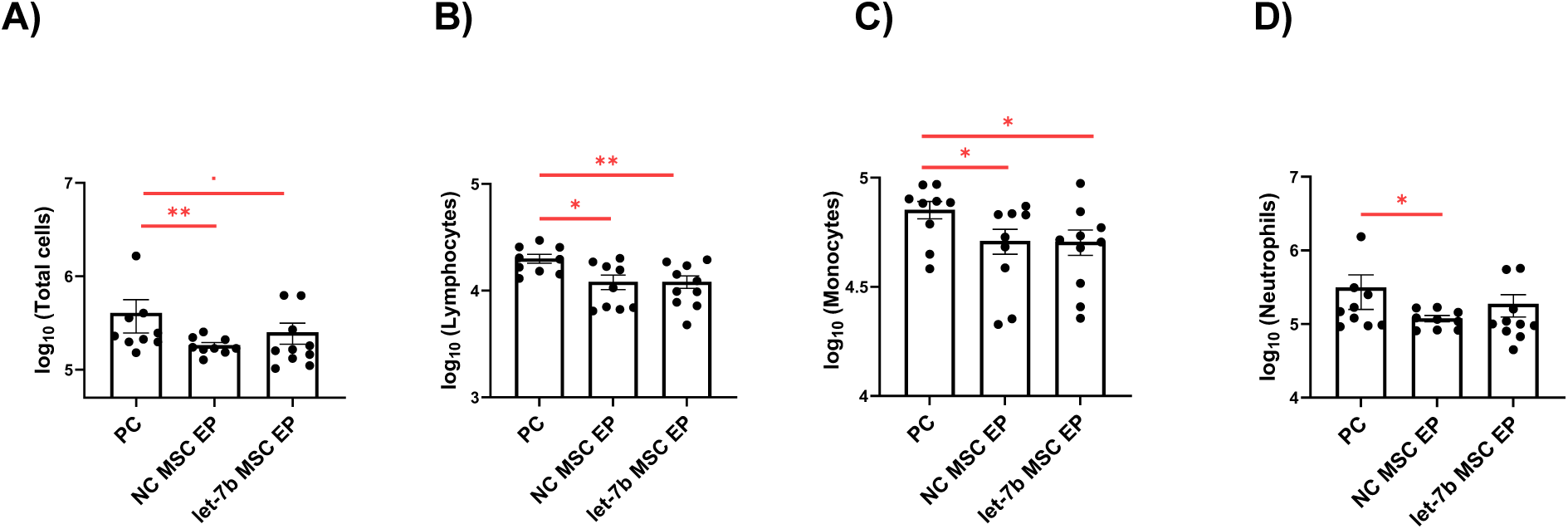
**A)** Total cell count in BALF was significantly reduced by NC MSC EPs (by 54.8%) (p = 0.00713) but was not significantly reduced by let-7b-5p MSC EPs (by 37.9%) (p = 0.06). **B)** MSC EPs loaded with negative control (NC) miRNA or let-7b-5p significantly decreased lymphocytes (39.4% by NC MSC EPs; 39.4% by let-7b-5p MSC EPs) in the BALF of CF mice as compared to PC [(p= 0.011 and 0.008 for NC (vs PC) and let-7b-5p (vs PC) respectively)]. **C)** MSC EPs loaded with negative control (NC) miRNA or let-7b-5p significantly decreased monocytes (27.9% by NC MSC EPs; 28.6% by let-7b-5p MSC EPs) (p-value = 0.03 and 0.02 for NC (vs PC) and let-7b-5p (vs PC) respectively). **D)** NC MSC EPs significantly reduced neutrophils (by 61.9%) (p = 0.018), whereas let-7b-5p MSC EPs tended to reduce neutrophils (p = 0.13), but the change was not significant (39.9%). The control group was exposed to process control (PC). Negative binomial generalized linear models with batch as a fixed effect were used to account for potential batch effects and test the statistical significance between the control and EP groups. Each data point represents one cystic fibrosis (CF) mouse. \**P* < 0.05, **P < 0.01; *n* = 9-10 mice/group; mean ± SEM depicted. Data were compiled from 3 independent experiments. PC = process control. A power analysis of the data (at p = 0.05 and power = 0.80) revealed that an additional 83 mice are needed to obtain statistically significant changes in neutrophils in let-7b-5p MSC EP compared to PC, making further mouse experiments impractical.

### Transfection of MSC with NC or let-7b-5p miRNAs altered the abundance of several miRNAs in EPs isolated from NC MSC and let-7b-5p MSC compared to EPs secreted by untreated MSC

To determine why let-7b-5p MSC EPs were not more effective than NC MSC EPs in altering cytokine secretion and immune cell counts, despite the well-known anti-bacterial and anti-inflammatory effect of let-7b-5p, studies were conducted to interrogate the miRNA content of MSC EPs, NC MSC EPs and let-7b-5p MSC EPs. miRNA-seq revealed that the transfection procedure altered the miRNA content of NC MSC EPs and let-7b-5p MSC EPs compared to untreated MSC EPs. The 20 miRNAs with the largest (10) and smallest (10) log_2_FC values for the three different preparations of MSC EPs are presented in Figure 6 [NC MSC EPs vs. EPs isolated from untransfected MSCs (**Figure 6A**); let-7b-5p MSC EPs vs. EPs isolated from untransfected MSCs (**Figure 6b**) and let-7b-5p MSC EPs vs. NC MSC EPs (**Figure 6C**)]. The data reveal that the transfection process changed the levels of miRNAs in NC MSC EPs and let-7b-5p MSC Eps compared to control MSC EPs. For instance, the abundance of some miRNAs either doubled (log_2_FC > 1) or halved (log_2_FC < -1) in the NC MSC EPs **(Figure 6A)** and let-7b-5p MSC EPs compared to the EPs isolated from untransfected MSCs **(Figure 6B)**.

**Figure 6:**
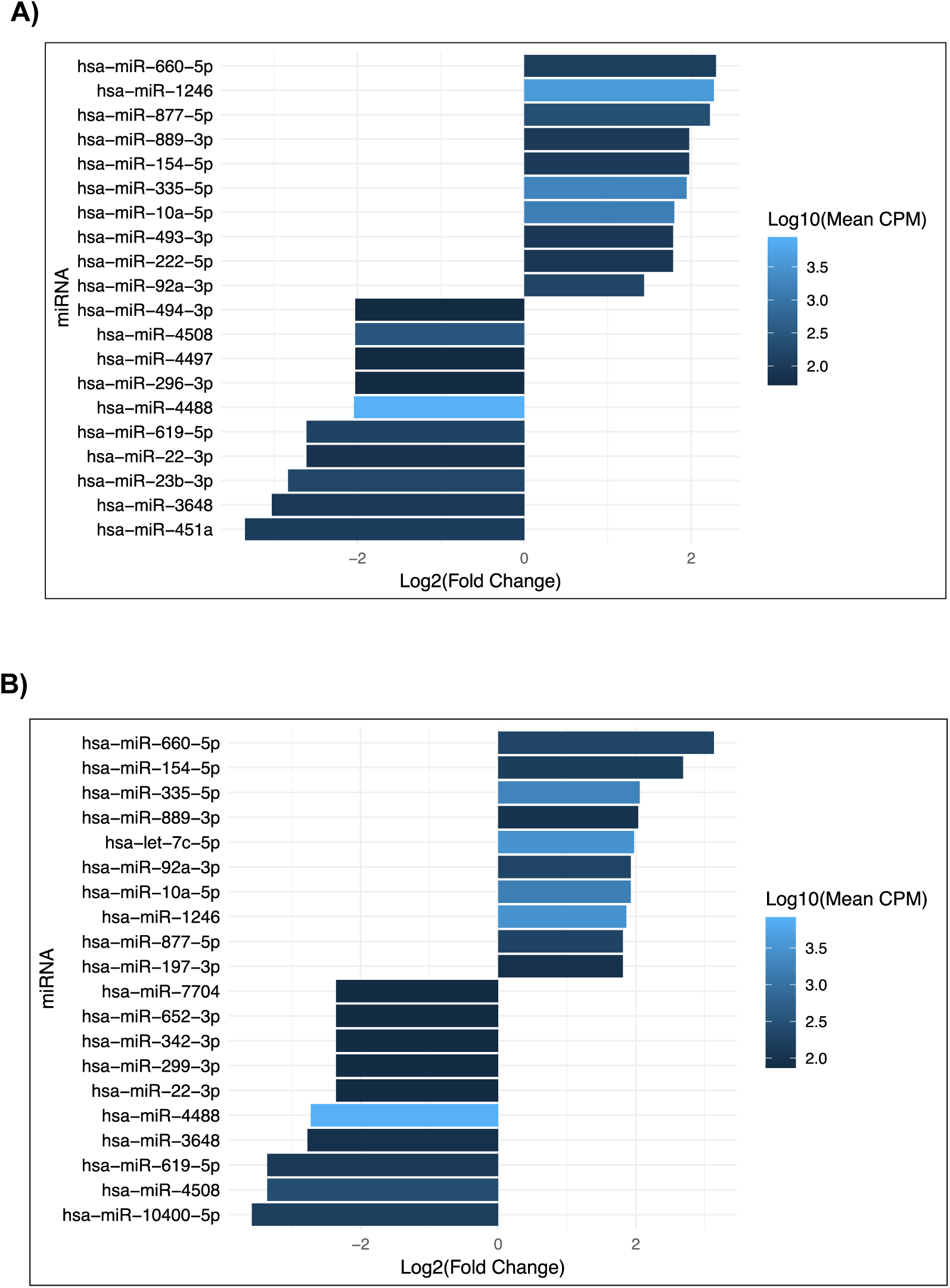
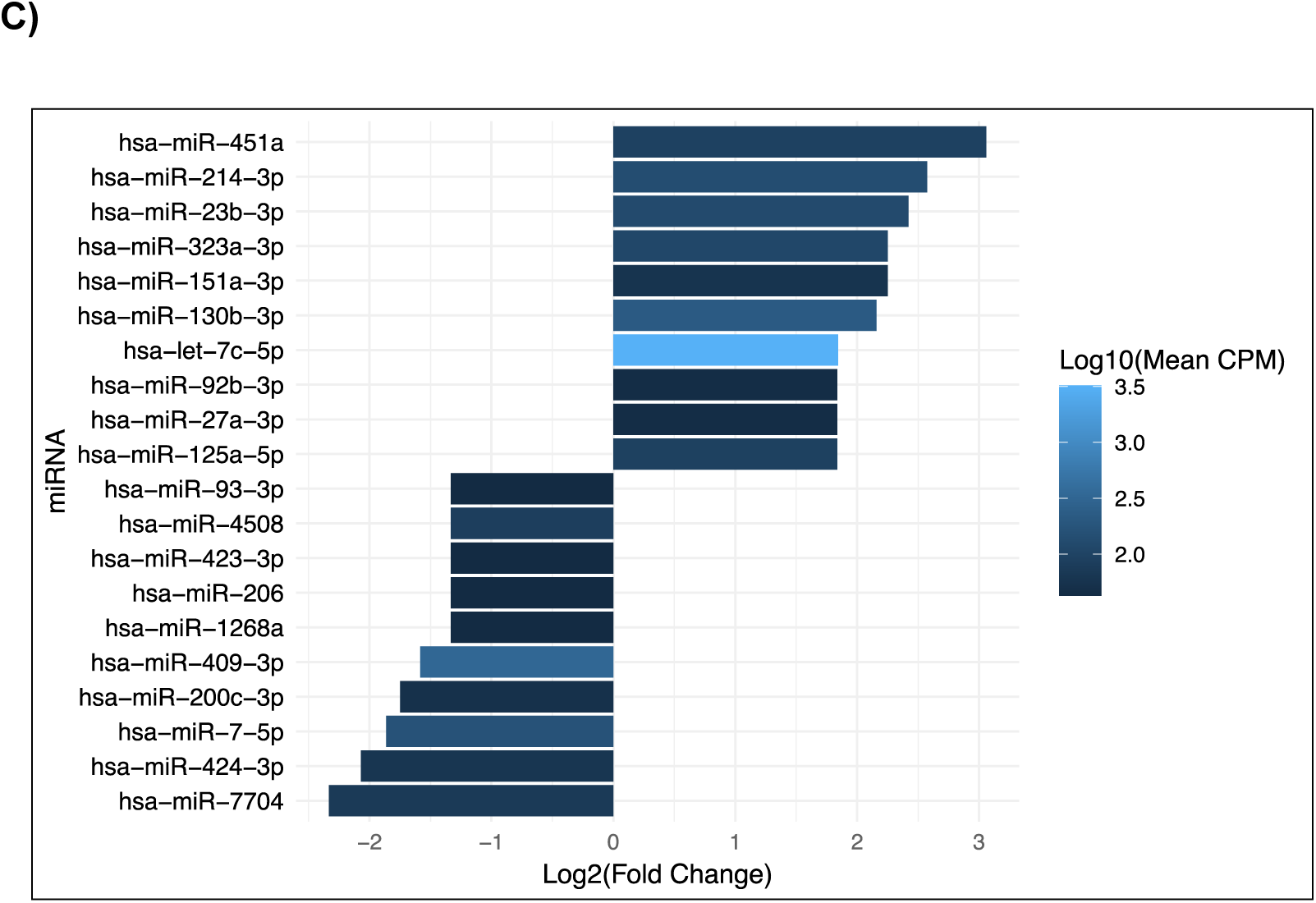
Comparison of miRNAs detected in: **A)** MSC EPs loaded with negative control (NC) miRNA vs. EPs isolated from untransfected MSCs; **B)** MSC EPs loaded with let-7b-5p miRNA vs. EPs isolated from untransfected MSCs and **C)** let-7b-5p loaded MSC EPs vs. NC loaded MSC EPs. The plots show the 20 miRNAs with the largest and smallest log_2_FC values for each comparison. Bar colors correspond to the mean counts per million calculated across the samples in each comparison. miRNA let-7b-5p was excluded from the analysis due to its large log_2_FC values that would dominate the graphs (see Figure 7 for let-7b-5p data).

Using the data in Figure 6 we identified 8 miRNAs whose abundance increased by over 100% and 5 miRNAs whose abundance decreased by more than 50% in both let-7b-5p MSC EPs and NC MSC EPs compared to the EPs isolated from untransfected MSCs. Notably, miR-877-5p, which is predicted to target many of the same genes in *Pseudomonas* as let-7b-5p, was elevated in both NC MSC EPs and let-7b-5p MSC EPs. miR-877-5p was increased to greater extent in NC MSC EPs than in let-7b-5p MSC EPs. The log_2_FC for miR-877-5p in NC MSC EPs vs untransfected MSC EPs was 2.22, whereas the same analysis for miR-877-5p in let-7b-5p MSC EPs vs untransfected MSC EPs was 1.81. miRNA-877-5p and let-7b-5p have greater than 50% overlap in their potential *P. aeruginosa* gene targets, as determined by the Rocket-miR application that we developed to predict druggable targets by miRNAs in 24 species of bacteria (41). This observation may explain why we saw a similar reduction in *P. aeruginosa* burden for both the let-7b-5p and NC MSC EP groups compared to PC in CF mice. We also explored the miRNA-seq data for other miRNAs upregulated in both NC and let-7b-5p EPs (compared to untreated EPs) that are predicted to target genes on *Pseudomonas* and ran them through our Rocket-miR (41). We found that two upregulated miRNAs, namely miR-154-5p and miR-92a-3p target the beta-lactam resistance pathway and biofilm formation pathway respectively in *P. aeruginosa.* Therefore, the antibacterial effect on *Pseudomonas* is likely a combination of the actions of several miRNAs beyond 877-5p and let-7b-5p, and several miRNA cargo is likely to reduce *Pseudomonas*. **Table 2** highlights the top 10 KEGG pathways of predicted miR-877-5p and let-7b-5p gene targets for *P. aeruginosa* and other CF pathogens. The other pathogens such as *Staphylococcus aureus*, *Streptococcus sanguinis*, and *Prevotella melaninogenica* were chosen because a data mining publication from our laboratory revealed that these bacteria accounted for 24% of the variability in lung function in a cohort of 167 pwCF (70). Pathways highlighted in red have greater than 50% of their genes predicted to be targets of let-7b-5p and miRNA-877-5p. These observations suggest that let-7b-5p would have important impacts on other bacteria, and that the upregulation of miRNA-877-5p in NC MSC EPs, induced by transfection, may account for the similar effects of NC MSC EPs and let-7b-5p MSC EPs.

**Table 2:** KEGG pathways in CF pathogens with the highest percentage of genes targeted by miR-877-5p, with pathways highlighted in red indicating those in which over 50% of genes are also targeted by let-7b-5p. Some of the genes in biofilm formation and antibiotic resistance pathways were identified previously by us as targeted by let-7b-5p and miRNA-877-5p using Rocket-miR (41).

| *Pseudomonas aeruginosa* | *Staphylococcus aureus* | *Streptococcus sanguinis* | *Prevotella melaninogenica* |
| --- | --- | --- | --- |
| Acarbose and validamycin biosynthesis<br>Naphthalene degradation<br>Degradation of flavonoids<br>Viral life cycle<br>Non-homologous end-joining<br>Pentose and glucuronate interconversions<br>Nucleotide excision repair<br>Starch and sucrose metabolism<br>RNA polymerase<br>Alanine, aspartate and glutamate metabolism | Xylene degradation<br>Benzoate degradation<br>Monobactam biosynthesis<br>Degradation of aromatic compounds<br>Valine, leucine and isoleucine degradation<br>Teichoic acid biosynthesis<br>Glutathione metabolism<br>Lysine degradation<br>Tryptophan metabolism<br>Fatty acid metabolism | Glycan degradation<br>Valine, leucine and isoleucine degradation<br>Penicillin and cephalosporin biosynthesis<br>Carbapenem biosynthesis<br>Sphingolipid metabolism<br>Flagellar assembly<br>Sulfur relay system<br>Selenocompound metabolism<br>Glutathione metabolism<br>Streptomycin biosynthesis | Degradation of flavonoids<br>Biosynthesis of various plant secondary metabolites<br>Beta-lactam resistance<br>Oxidative phosphorylation<br>Nucleotide excision repair<br>Ubiquinone and other terpenoid quinone biosynthesis<br>Mismatch repair<br>Histidine metabolism<br>Flagellar assembly<br>Peptidoglycan biosynthesis |

Importantly, using IntaRNA and the DAVID functional annotation tool, we found that 16 miRNAs were differentially expressed and predicted to reduce inflammation in mice. For instance, miRNA-206 and miRNA-424-3p increased in NC EPs relative to let-7b-5p MSC EPs. Both miRNAs are predicted to inhibit genes involved in PI3K-Akt and MAPK proinflammatory signaling, respectively. miRNA-451a and miRNA-125a-5p increased in let-7b-5p MSC EPs compared to NC MSC EPs and target genes involved in B-cell activation and NF-κB signaling, respectively **(Table 3)**. Thus, both NC MSC EPs and let-7b-5p EPs contain different sets of anti-inflammatory miRNAs possibly providing an explanation as to why both sets of MSC EPs reduced inflammation.

**Table 3:**
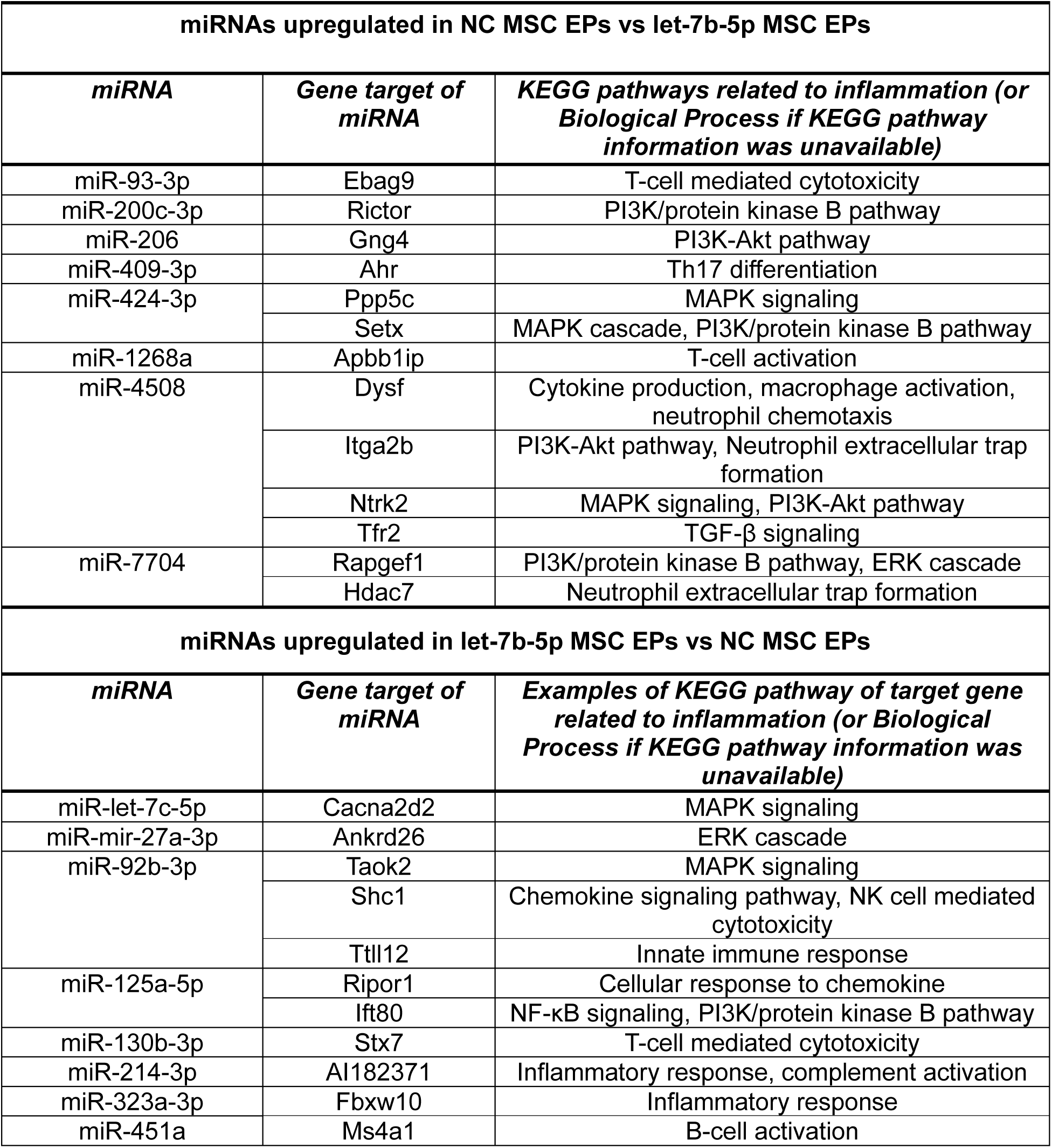
miRNAs in NC MSC EPs and let-7b-5p MSC EPs are predicted to have differential effects on several inflammatory pathways in mice.

Given our interest in using MSC EPs as a potential therapy, we investigated potential miRNA targets in human genes involved in inflammation. Our analysis revealed that several miRNAs differentially expressed in MSC EPs containing NC or let-7b-5p are also predicted to target key inflammatory pathways in humans. For example, miRNAs miR-206 and miR-93-3p enriched in NC EPs are predicted to suppress genes associated with NF-κB and JAK-STAT signaling. miRNAs miR-27a-3p and miR-92b-3p, which are more abundant in let-7b-5p MSC EPs are predicted to suppress genes involved in PI3K and MAPK pathways **(Table 4)**. This difference in the miRNA makeup may explain why NC MSC EPs and let-7b-5p MSC EPs have slightly different effects on the secretion of some cytokines and immune cells in BALF, and is a topic for further investigation. In summary, the miRNA analysis identified several miRNAs that may be responsible for the observed reduction in *P. aeruginosa* burden as well as anti-inflammatory effects observed in CF mice treated with NC MSC EPs and let-7b-5p MSC-EPs compared to untransfected MSC EPs.

**Table 4:** miRNAs in NC MSC EPs and let-7b-5p MSC EPs that are predicted to have differential effects on several inflammatory KEGG pathways in humans.

| **miRNAs upregulated in NC MSC EPs vs let-7b-5p MSC EPs** |  |  |
| --- | --- | --- |
| ***miRNA*** | ***Gene target of miRNA*** | ***KEGG pathways of target gene related to inflammation (or Biological Process if KEGG pathway information was unavailable)*** |
| miR-93-3p | MYO18A | NF-κB transduction, macrophage activation |
| miR-206 | ZFP91 | NF-κB transduction |
|  | SOCS1 | JAK STAT signaling |
| miR-409-3p | JUN | MAPK signaling, TLR signaling, IL-17 signaling, TNF signaling |
| miR-423-3p | CCDC88A | PI3K signaling |
|  | MRAS | MAPK signaling |
| miR-4508 | CSPG4 | MAPK signaling |
| **miRNAs upregulated in let-7b-5p MSC EPs vs NC MSC EPs** |  |  |
| ***miRNA*** | ***Gene target of miRNA*** | ***KEGG pathway of target gene related to inflammation (or Biological Process if KEGG pathway information was unavailable)*** |
| miR-let-7c-5p | CMKLR1 | Inflammatory response, macrophage chemotaxis, complement receptor mediated signaling |
|  | LTBR4 | Inflammatory response, immune response |
| miR-23b-3p | TNFSF8 | Cytokine-cytokine receptor interaction |
|  | DLL4 | T-cell differentiation |
| miR-27a-3p | PIK3C2A | PI3K pathway |
| miR-92b-3p | TAOK2 | MAPK signaling |
|  | CDKN1A | JAK STAT signaling, PI3K-Akt pathway |
| miR-125a-5p | ZBTB33 | Cytokine production regulation |
| miR-130b-3p | RGMB | TGF-β signaling |
| miR-214-3p | HRNA4 | B-cell activation |
|  | STAB1 | Inflammatory response |
| miR-323a-3p | CFLAR | NF-κB signaling, TNF signaling |
| miR-451a | NPNT | ERK cascade |

In addition, smRNA-seq validated the qPCR experiments (Figure 1B) demonstrating that transfection of MSC with let-7b-5p increased its abundance compared to NC MSC EPs and untransfected MSC **(Figure 7)**.

**Figure 7:**
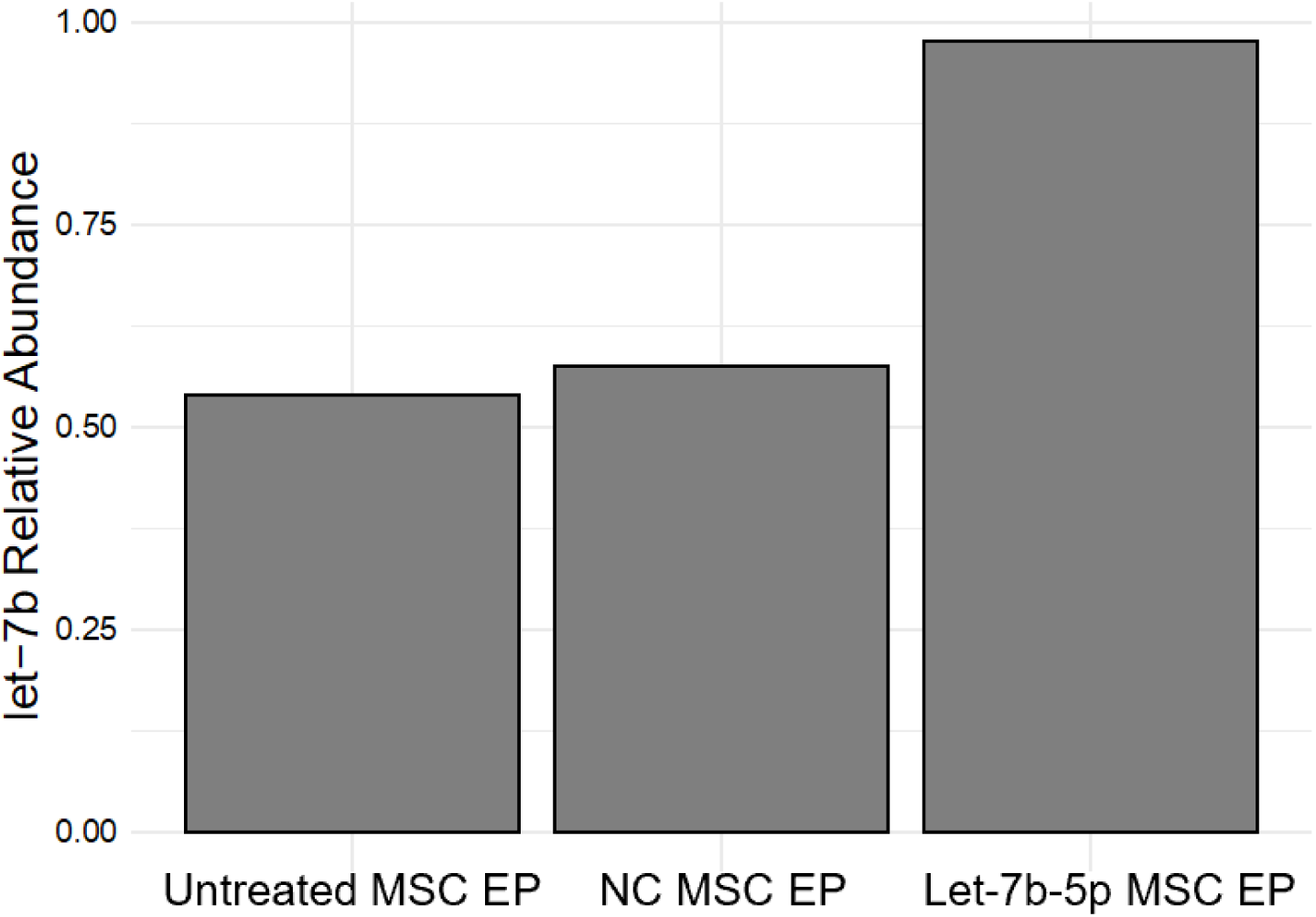
Relative abundance of let-7b-5p miRNA transcripts in EPs isolated from untransfected MSCs (Untreated MSC EP), EPs isolated from MSCs transfected with a NC miRNA (NC MSC EP) and EPs isolated from MSCs transfected with let-7b-5p (let-7b-5p MSC EP).

## DISCUSSION

Our mouse model of infection demonstrated that MSC EPs effectively reduced the *P. aeruginosa* burden and immune cell recruitment in the lungs while shifting the cytokine profile toward a more anti-inflammatory/pro-resolving landscape, even in the absence of antibiotics. MSC EPs increased IL-10, an anti-inflammatory and pro-resolving cytokine in BALF and decreased serum RANTES, a chemoattractant for immune cells. In addition, MSC EPs reduced monocytes, neutrophils and lymphocytes in BALF of CF mice. Our study is in general agreement with the studies of Shi et al (31) and extend the significance of MSC EPs to a frequently utilized CF mouse model of acute *P. aeruginosa* infection, which manifests excessive inflammation and a reduced ability to eliminate infections compared to non-CF mice (71). Shi et al. showed that MSC EPs significantly reduced mortality, neutrophil influx, proinflammatory factors, and bacterial burden in the BALF of non-CF mice with multidrug-resistant *Pseudomonas aeruginosa* pneumonia (31). Park et al. found that intravenous administration of MSC EPs effectively reduced bacterial burden in ex-vivo human lung tissue with severe bacterial pneumonia (30). The present study is also consistent with a previous report in which we demonstrated that EPs secreted by primary human bronchial epithelial cells endogenously containing let-7b-5p suppressed *P. aeruginosa* burden, reduced immune cells in BALF and decreased the proinflammatory response to *P. aeruginosa* even in the absence of antibiotics (42).

The inability of MSC EPs containing let-7b-5p to reduce bacterial burden compared to MSC Eps + NC was not surprising given our earlier *in vitro* study demonstrating that let-7b-5p in combination with sub minimum inhibitory concentrations of either aztreonam or carbenicillin was required to reduce *P. aeruginosa* (38). The combination of let-7b-5p and aztreonam was required to reduce *P. aeruginosa* due to the mechanism of action of let-7b-5p, which is to reduce aztreonam antibiotic efflux pumps, rendering aztreonam more effective in killing *P. aeruginosa* (38). The mechanism of action of EPs is usually attributed to the release of their cargo into recipient cells. We have shown that HBEC EPs transfer the miRNA let-7b-5p to *Pseudomonas*, and blocking the miRNA via an anti-sense nucleotide to let-7b-5p abolished the ability of let-7b-5p plus aztreonam to reduce *P. aeruginosa* CFUs, by blocking the inhibition of genes involved in biofilm formation and B-lactam resistance (38). Direct expression of let-7b-5p in *Pseudomonas* had a similar effect, indicating that let-7b-5p in combination with aztreonam reduces *Pseudomonas* CFUs. Additional studies are required to evaluate the ability of MSC EPs and aztreonam together in mice to reduce *P. aeruginosa* burden and reduce inflammation in CF mice.

Studies were also conducted to determine why NC MSC EPs and let-7b-5p MSC EPs had many similar effects in mice. Analysis of the miRNA content of the MSC EPs revealed that transfection of a control miRNA as well as let-7b-5p in MSC changed the abundance of other miRNAs in EPs compared to untreated MSC. Most notably, transfection with the control miRNA increased the expression of miRNA-877-5p, which has similar predicted gene targets in *P. aeruginosa* as let-7b-5p. miR-877-5p was increased to a greater extent in NC MSC EPs (log_2_FC of 2.22 vs untreated) than in let-7b-5p MSC EPs (log_2_FC of 1.81 vs untreated). Examining the most prevalent miRNAs with predicted antibacterial properties, we propose that the effects of miRNA-877-5p (and possibly other miRNAs or other components in NC MSC EPs) may mediate the reduction of *P. aeruginosa* in mice exposed to NC MSC EPs, whereas let-7b-5p (and possibly other miRNAs or other components in let-7b-5p MSC EPs) reduced *P. aeruginosa* in mice exposed to let-7b-5p MSC EPs.

Elucidating the effects of the relative changes in numerous miRNAs, and possibly other components, in the MSC EPs on *P. aeruginosa,* immune cells and cytokine abundance is beyond the goals of this study and will require an intensive effort to unravel. However, analysis with our Rocket-miR (41) program found that two upregulated miRNAs in the two sets of EPs, namely miR-154-5p and miR-92a-3p target the beta-lactam resistance pathway and biofilm formation pathway respectively in *P. aeruginosa.* Therefore, the antibacterial effect is likely a combination of the actions of several miRNAs beyond 877-5p and let-7b-5p, and possibly other miRNA cargo in EPs. Additionally, it is possible that transfection can alter not only miRNA cargo but also other EP components, such as lipids and proteins, which have been shown to play a key role in mediating antibacterial effects (72). Though beyond the scope of the present study, future investigations will be needed to dissect the contribution of these non-miRNA EP components to the observed antibacterial responses.

Our study has several notable strengths. First, the CF mouse model has been used extensively to study *P. aeruginosa* lung infections, thus, it is a well characterized and established model (56). Second, developing effective anti-Pseudomonal therapies is essential for all people with antibiotic resistant infections by *Pseudomonas,* including those with COPD and pneumonia (73). In addition, the Rocket-miR program also predicts that several miRNAs may have bactericidal effects on a number of other pathogens. The ability of MSC EPs to reduce bacterial burden and inflammation underscores their potential as a therapeutic option. MSC EPs are particularly compelling due to their dual mechanism of action: potent anti-bacterial and anti-inflammatory properties, the latter of which could help prevent infection-induced lung damage. Additionally, MSC EPs are known for their intrinsic pro-resolving, safe, and non-toxic characteristics, distinguishing them from conventional anti-inflammatory drugs that often have severe side effects. A Phase I clinical trial demonstrated the safety and non-toxic effects of parent MSCs in adults with CF (21), providing a strong foundation for exploring MSC EPs in pwCF. Our focus on the efficacy of engineered MSC EPs in mitigating bacterial burden, particularly against clinical mucoid isolates of *Pseudomonas aeruginosa*, is an important step forward. Third, the use of process control (unconditioned media subjected to EP isolation process) instead of saline or PBS as a control has several advantages, including minimizing potential factors in culture media or contaminants that may be added inadvertently in the isolation process that may affect infection and inflammation. Finally, MSC EPs may have beneficial effects, including reducing inflammation, in several other disease models including Chronic Obstructive Pulmonary Disease (COPD), COVID-19, asthma, acute respiratory distress syndrome (ARDS), and idiopathic pulmonary fibrosis (IPF) (74), all via transfer of different miRNA/protein factors into target cells. Thus, we predict that the use of MSC EPs engineered to contain anti-bacterial and anti-inflammatory miRNAs will be beneficial to numerous disease states characterized by antibiotic resistant bacteria and excessive inflammation. Moreover, we anticipate that our Rocket-miR program will facilitate identification of miRNAs that target genes involved in all antibiotic-resistant infections.

Despite its strengths, our study has some limitations. First, the agarose bead model in CF mice allows for only short-term infection studies (up to 10 days) and does not fully replicate the chronic bacterial infections seen in pwCF, although the bactericidal effects of antibiotics in mice frequently translates to studies in humans (6, 42, 56). Moreover, there was considerable variation in the response of mice in all experimental sets, which is not uncommon in experiments in mice, but still presents difficulties in detecting small changes in responses. Another limitation of this study is the simultaneous administration of MSC EPs and bacteria. Future studies will explore staggered treatments, where bacteria are introduced first, followed by MSC EPs. This approach would more closely mimic clinical scenarios where treatment is initiated after the onset of infection. In addition, although in a few assays there appeared to be an effect of let-7b-5p MSC EPs compared to PC, the changes were not significant. We considered conducting additional studies to determine if the trend would be significant, however, a power analysis indicated that an additional 48 - 140 mice would be needed, which is not realistic. Finally, while this study focused on *Pseudomonas aeruginosa*, which colonizes approximately 50% of adults with CF, the CF lung microbiome is typically polymicrobial, including pathogens like *S. aureus*, *S. sanguinis*, *P. melaninogenica*, *Haemophilus influenzae*, and *Burkholderia cenocepacia* and the presence of other bacteria may affect the response of *P. aeruginosa* to MSC EPs, since the presence of other bacteria affect gene expression and antibiotic resistance of *P. aeruginosa* (75, 76). Our program Rocket-miR (41) predicts that let-7b-5p targets genes in other bacteria common in lung infections, including *P. melaninogenica*, *S. aureus*, and *S. sanguinis*. Future studies should explore the effects of let-7b-5p MSC EPs on polymicrobial infections to broaden the therapeutic scope. In addition, Rocket-miR also identified several other miRNAs that are predicted to target key genes in 24 bacterial genera, including all 6 ESKAPE (*Enterococcus faecium*, *Staphylococcus aureus*, *Klebsiella pneumoniae*, *Acinetobacter baumannii*, *Pseudomonas aeruginosa* and *Enterobacter* spp.) pathogens, which are highly virulent and often antibiotic resistant.

In conclusion, our data demonstrates that MSC EPs reduce *P. aeruginosa* infection, immune cells and inflammation in our CF mouse model of infection. We hypothesize three potential mechanisms by which MSC EPs reduce bacterial burden and inflammation, none mutually exclusive. First, MSC EPs exhibit direct bactericidal activity, as shown in our previous studies (19). Second, MSC EPs may enhance phagocytosis and bacterial killing by CF macrophages and other immune cells, thereby reducing bacterial load. Third, MSC EPs may increase the secretion of antimicrobial peptides by airway epithelial cells. Future experiments will test these mechanisms to better understand how MSC EPs function *in vivo*. This knowledge will be instrumental in developing MSC EPs as dual-action anti-infective and anti-inflammatory therapy for all people with chronic infection and inflammation.

## ACKNOWLEDGEMENTS

This study was supported by the Cystic Fibrosis Foundation (STANTO19G0, STANTO20P0, STANTO23R0, and STANTO19R0) and the National Institutes of Health (P30-DK117469 and R01HL151385) to BAS and NIH F31HL172440 to LAC. This work was supported by CFF RDP R447-CR11 and CFF HODGES19R1 (Phenotyping Core). We thank RoosterBio for analysis of the MSC EPs (Table 1 and Figure 1A). smRNA-seq was carried out in the Genomics and Molecular Biology Shared Resource (RRID:SCR_021293) at Dartmouth which is supported by NCI Cancer Center Support Grant 5P30CA023108 and NIH S10 (1S10OD030242) awards. Support for RNA-seq data analysis was provided through Dartmouth Cancer Center Bioinformatics & Biostatistics Shared Resource (BBSR) supported by NCI award P30CA023108 and Dartmouth’s Center for Quantitative Biology (CQB) Genomics Data Sciences Core (GDSC) supported by NIH award 3P20GM130454. Mass Spectrometry analyses were performed by the Mass Spectrometry Technology Access Center at the McDonnell Genome Institute (MTAC@MGI) at Washington University School of Medicine, supported by the Diabetes Research Center/NIH grant P30 DK020579, Institute of Clinical and Translational Sciences/NCATS CTSA award UL1 TR002345, and Siteman Cancer Center/NCI CCSG grant P30 CA091842.

## DATA AVAILABILITY

Data will be made available upon reasonable request. EP sequencing data has been uploaded to NCBI GEO (Accession number: GSE290294). Detailed methods has been uploaded to EV-track (77) to enhance rigor and reproducibility. The mass spectrometry proteomics data have been deposited to the ProteomeXchange Consortium via the PRIDE (78) partner repository with the dataset identifier PXD070855.

## DISCLAIMERS

The funders had no role in study design, data collection and analysis, publication decisions, or manuscript preparation.

## AUTHOR CONTRIBUTIONS

S.S., R.B., Z.F., M.J.W., D.J.W., T.L.B., T.J.K., and B.A.S. conceived and designed research; S.S., R.B., Z.F., C.R., B.V., B.K.C., and Y.A.G performed experiments; S.S., L.A.C., L.T., T.H.H, B.K.C., and Y.A.G analyzed data; S.S., R.B., Z.F., L.T., T.J.K., and B.A.S. interpreted results of experiments; S.S., L.A.C., L.T., and B.V. prepared figures; S.S. and B.A.S. drafted the manuscript; all authors edited and revised the manuscript; all authors approved the final version of the manuscript.

## REFERENCES

1. Karruli A, Catalini C, D’Amore C, Foglia F, Mari F, Harxhi A, Galdiero M, Durante-Mangoni E. Evidence-Based Treatment of Pseudomonas aeruginosa Infections: A Critical Reappraisal. Antibiotics (Basel) 12: 399, 2023. doi: 10.3390/antibiotics12020399.

2. Nichols DP, Paynter AC, Heltshe SL, Donaldson SH, Frederick CA, Freedman SD, Gelfond D, Hoffman LR, Kelly A, Narkewicz MR, Pittman JE, Ratjen F, Rosenfeld M, Sagel SD, Schwarzenberg SJ, Singh PK, Solomon GM, Stalvey MS, Clancy JP, Kirby S, Van Dalfsen JM, Kloster MH, Rowe SM, PROMISE Study group. Clinical Effectiveness of Elexacaftor/Tezacaftor/Ivacaftor in People with Cystic Fibrosis: A Clinical Trial. Am J Respir Crit Care Med 205: 529–539, 2022. doi: 10.1164/rccm.202108-1986OC.

3. Allen L, Allen L, Carr SB, Davies G, Downey D, Egan M, Forton JT, Gray R, Haworth C, Horsley A, Smyth AR, Southern KW, Davies JC. Future therapies for cystic fibrosis. Nat Commun 14: 693, 2023. doi: 10.1038/s41467-023-36244-2.

4. Krause KM, Haglund CM, Hebner C, Serio AW, Lee G, Nieto V, Cohen F, Kane TR, Machajewski TD, Hildebrandt D, Pillar C, Thwaites M, Hall D, Miesel L, Hackel M, Burek A, Andrews LD, Armstrong E, Swem L, Jubb A, Cirz RT. Potent LpxC Inhibitors with In Vitro Activity against Multidrug-Resistant Pseudomonas aeruginosa. Antimicrob Agents Chemother 63: e00977–19, 2019. doi: 10.1128/AAC.00977-19.

5. Taccetti G, Francalanci M, Pizzamiglio G, Messore B, Carnovale V, Cimino G, Cipolli M. Cystic Fibrosis: Recent Insights into Inhaled Antibiotic Treatment and Future Perspectives. Antibiotics (Basel) 10: 338, 2021. doi: 10.3390/antibiotics10030338.

6. Roesch EA, Nichols DP, Chmiel JF. Inflammation in cystic fibrosis: An update. Pediatr Pulmonol 53: S30–S50, 2018. doi: 10.1002/ppul.24129.

7. Caverly LJ, Riquelme SA, Hisert KB. The Impact of Highly Effective Modulator Therapy on Cystic Fibrosis Microbiology and Inflammation. Clin Chest Med 43: 647–665, 2022. doi: 10.1016/j.ccm.2022.06.007.

8. Keown K, Brown R, Doherty DF, Houston C, McKelvey MC, Creane S, Linden D, McAuley DF, Kidney JC, Weldon S, Downey DG, Taggart CC. Airway Inflammation and Host Responses in the Era of CFTR Modulators. Int J Mol Sci 21: 6379, 2020. doi: 10.3390/ijms21176379.

9. Chmiel JF, Konstan MW, Elborn JS. Antibiotic and anti-inflammatory therapies for cystic fibrosis. Cold Spring Harb Perspect Med 3: a009779, 2013. doi: 10.1101/cshperspect.a009779.

10. Antimicrobial Resistance Collaborators. Global burden of bacterial antimicrobial resistance in 2019: a systematic analysis. Lancet 399: 629–655, 2022. doi: 10.1016/S0140-6736(21)02724-0.

11. Pittenger MF, Discher DE, Péault BM, Phinney DG, Hare JM, Caplan AI. Mesenchymal stem cell perspective: cell biology to clinical progress. NPJ Regen Med 4: 22, 2019. doi: 10.1038/s41536-019-0083-6.

12. Bonfield TL, Sutton MT, Fletcher DR, Folz MA, Ragavapuram V, Somoza RA, Caplan AI. Donor-defined mesenchymal stem cell antimicrobial potency against nontuberculous mycobacterium. Stem Cells Transl Med 10: 1202–1216, 2021. doi: 10.1002/sctm.20-0521.

13. Rolandsson Enes S, Hampton TH, Barua J, McKenna DH, Dos Santos CC, Amiel E, Ashare A, Liu KD, Krasnodembskaya AD, English K, Stanton BA, Rocco PRM, Matthay MA, Weiss DJ. Healthy versus inflamed lung environments differentially affect mesenchymal stromal cells. Eur Respir J 58: 2004149, 2021. doi: 10.1183/13993003.04149-2020.

14. O’Kane CM, Matthay MA. Understanding the Role of Mesenchymal Stromal Cells in Treating COVID-19 Acute Respiratory Distress Syndrome. Am J Respir Crit Care Med 207: 231–233, [date unknown]. doi: 10.1164/rccm.202209-1838ED.

15. Le Blanc K, Tammik C, Rosendahl K, Zetterberg E, Ringdén O. HLA expression and immunologic properties of differentiated and undifferentiated mesenchymal stem cells. Exp Hematol 31: 890–896, 2003. doi: 10.1016/s0301-472x(03)00110-3.

16. Wu X, Jiang J, Gu Z, Zhang J, Chen Y, Liu X. Mesenchymal stromal cell therapies: immunomodulatory properties and clinical progress. Stem Cell Res Ther 11: 345, 2020. doi: 10.1186/s13287-020-01855-9.

17. Babajani A, Soltani P, Jamshidi E, Farjoo MH, Niknejad H. Recent Advances on Drug-Loaded Mesenchymal Stem Cells With Anti-neoplastic Agents for Targeted Treatment of Cancer. Front Bioeng Biotechnol 8: 748, 2020. doi: 10.3389/fbioe.2020.00748.

18. Zhuang W-Z, Lin Y-H, Su L-J, Wu M-S, Jeng H-Y, Chang H-C, Huang Y-H, Ling T-Y. Mesenchymal stem/stromal cell-based therapy: mechanism, systemic safety and biodistribution for precision clinical applications. J Biomed Sci 28: 28, 2021. doi: 10.1186/s12929-021-00725-7.

19. Sutton MT, Fletcher D, Ghosh SK, Weinberg A, van Heeckeren R, Kaur S, Sadeghi Z, Hijaz A, Reese J, Lazarus HM, Lennon DP, Caplan AI, Bonfield TL. Antimicrobial Properties of Mesenchymal Stem Cells: Therapeutic Potential for Cystic Fibrosis Infection, and Treatment. Stem Cells Int 2016: 5303048, 2016. doi: 10.1155/2016/5303048.

20. Center for Drug Evaluation and Research. FDA approves remestemcel-L-rknd for steroid-refractory acute graft versus host disease in pediatric patients [Online]. FDA FDA: 2024. https://www.fda.gov/drugs/resources-information-approved-drugs/fda-approves-remestemcel-l-rknd-steroid-refractory-acute-graft-versus-host-disease-pediatri [25Mar.2025].

21. Roesch EA, Bonfield TL, Lazarus HM, Reese J, Hilliard K, Hilliard J, Khan U, Heltshe S, Gluvna A, Dasenbrook E, Caplan AI, Chmiel JF. A phase I study assessing the safety and tolerability of allogeneic mesenchymal stem cell infusion in adults with cystic fibrosis. J Cyst Fibros 22: 407–413, 2023. doi: 10.1016/j.jcf.2022.12.001.

22. Börger V, Bremer M, Ferrer-Tur R, Gockeln L, Stambouli O, Becic A, Giebel B. Mesenchymal Stem/Stromal Cell-Derived Extracellular Vesicles and Their Potential as Novel Immunomodulatory Therapeutic Agents. Int J Mol Sci 18: 1450, 2017. doi: 10.3390/ijms18071450.

23. Lai RC, Arslan F, Lee MM, Sze NSK, Choo A, Chen TS, Salto-Tellez M, Timmers L, Lee CN, El Oakley RM, Pasterkamp G, de Kleijn DPV, Lim SK. Exosome secreted by MSC reduces myocardial ischemia/reperfusion injury. Stem Cell Res 4: 214–222, 2010. doi: 10.1016/j.scr.2009.12.003.

24. Bruno S, Grange C, Deregibus MC, Calogero RA, Saviozzi S, Collino F, Morando L, Busca A, Falda M, Bussolati B, Tetta C, Camussi G. Mesenchymal stem cell-derived microvesicles protect against acute tubular injury. J Am Soc Nephrol 20: 1053–1067, 2009. doi: 10.1681/ASN.2008070798.

25. Abreu SC, Lopes-Pacheco M, Weiss DJ, Rocco PRM. Mesenchymal Stromal Cell-Derived Extracellular Vesicles in Lung Diseases: Current Status and Perspectives. Front Cell Dev Biol 9: 600711, 2021. doi: 10.3389/fcell.2021.600711.

26. Williams T, Salmanian G, Burns M, Maldonado V, Smith E, Porter RM, Song YH, Samsonraj RM. Versatility of mesenchymal stem cell-derived extracellular vesicles in tissue repair and regenerative applications. Biochimie 207: 33–48, 2023. doi: 10.1016/j.biochi.2022.11.011.

27. Kou M, Huang L, Yang J, Chiang Z, Chen S, Liu J, Guo L, Zhang X, Zhou X, Xu X, Yan X, Wang Y, Zhang J, Xu A, Tse H-F, Lian Q. Mesenchymal stem cell-derived extracellular vesicles for immunomodulation and regeneration: a next generation therapeutic tool? Cell Death Dis 13: 580, 2022. doi: 10.1038/s41419-022-05034-x.

28. Aguiar Koga BA, Fernandes LA, Fratini P, Sogayar MC, Carreira ACO. Role of MSC-derived small extracellular vesicles in tissue repair and regeneration. Front Cell Dev Biol 10: 1047094, 2022. doi: 10.3389/fcell.2022.1047094.

29. Guo H, Su Y, Deng F. Effects of Mesenchymal Stromal Cell-Derived Extracellular Vesicles in Lung Diseases: Current Status and Future Perspectives. Stem Cell Rev Rep 17: 440–458, 2021. doi: 10.1007/s12015-020-10085-8.

30. Park J, Kim S, Lim H, Liu A, Hu S, Lee J, Zhuo H, Hao Q, Matthay MA, Lee J-W. Therapeutic effects of human mesenchymal stem cell microvesicles in an ex vivo perfused human lung injured with severe E. coli pneumonia. Thorax 74: 43–50, 2019. doi: 10.1136/thoraxjnl-2018-211576.

31. Shi M-M, Zhu Y-G, Yan J-Y, Rouby J-J, Summah H, Monsel A, Qu J-M. Role of miR-466 in mesenchymal stromal cell derived extracellular vesicles treating inoculation pneumonia caused by multidrug-resistant Pseudomonas aeruginosa. Clin Transl Med 11: e287, 2021. doi: 10.1002/ctm2.287.

32. Li X, Wang S, Zhu R, Li H, Han Q, Zhao RC. Lung tumor exosomes induce a pro-inflammatory phenotype in mesenchymal stem cells via NFκB-TLR signaling pathway. J Hematol Oncol 9: 42, 2016. doi: 10.1186/s13045-016-0269-y.

33. Suk JS, Lai SK, Wang Y-Y, Ensign LM, Zeitlin PL, Boyle MP, Hanes J. The penetration of fresh undiluted sputum expectorated by cystic fibrosis patients by non-adhesive polymer nanoparticles. Biomaterials 30: 2591–2597, 2009. doi: 10.1016/j.biomaterials.2008.12.076.

34. Long Q, Upadhya D, Hattiangady B, Kim D-K, An SY, Shuai B, Prockop DJ, Shetty AK. Intranasal MSC-derived A1-exosomes ease inflammation, and prevent abnormal neurogenesis and memory dysfunction after status epilepticus. Proc Natl Acad Sci U S A 114: E3536–E3545, 2017. doi: 10.1073/pnas.1703920114.

35. Reis M, Ogonek J, Qesari M, Borges NM, Nicholson L, Preußner L, Dickinson AM, Wang X-N, Weissinger EM, Richter A. Recent Developments in Cellular Immunotherapy for HSCT-Associated Complications. Front Immunol 7: 500, 2016. doi: 10.3389/fimmu.2016.00500.

36. Nassar W, El-Ansary M, Sabry D, Mostafa MA, Fayad T, Kotb E, Temraz M, Saad A-N, Essa W, Adel H. Erratum to: Umbilical cord mesenchymal stem cells derived extracellular vesicles can safely ameliorate the progression of chronic kidney diseases. Biomater Res 21: 3, 2017. doi: 10.1186/s40824-017-0089-3.

37. Kordelas L, Rebmann V, Ludwig A-K, Radtke S, Ruesing J, Doeppner TR, Epple M, Horn PA, Beelen DW, Giebel B. MSC-derived exosomes: a novel tool to treat therapy-refractory graft-versus-host disease. Leukemia 28: 970–973, 2014. doi: 10.1038/leu.2014.41.

38. Koeppen K, Nymon A, Barnaby R, Bashor L, Li Z, Hampton TH, Liefeld AE, Kolling FW, LaCroix IS, Gerber SA, Hogan DA, Kasetty S, Nadell CD, Stanton BA. Let-7b-5p in vesicles secreted by human airway cells reduces biofilm formation and increases antibiotic sensitivity of P. aeruginosa. Proc Natl Acad Sci U S A 118: e2105370118, 2021. doi: 10.1073/pnas.2105370118.

39. Qiu G, Zheng G, Ge M, Wang J, Huang R, Shu Q, Xu J. Mesenchymal stem cell-derived extracellular vesicles affect disease outcomes via transfer of microRNAs. Stem Cell Res Ther 9: 320, 2018. doi: 10.1186/s13287-018-1069-9.

40. Rolandsson Enes S, Dzneladze I, Hampton TH, Neff SL, Asarian L, Barua J, Tertel T, Giebel B, Pereyra N, McKenna DH, Hu P, Acton E, Ashare A, Liu KD, Krasnodembskaya AD, English K, Stanton BA, Rocco PRM, Matthay MA, dos Santos CC, Weiss DJ. Acute respiratory distress vs healthy lung environments differently affect mesenchymal stromal cell extracellular vesicle miRNAs. Cytotherapy 27: 581–596, 2025. doi: 10.1016/j.jcyt.2025.01.006.

41. Neff SL, Hampton TH, Koeppen K, Sarkar S, Latario CJ, Ross BD, Stanton BA. Rocket-miR, a translational launchpad for miRNA-based antimicrobial drug development. mSystems 8: e0065323, 2023. doi: 10.1128/msystems.00653-23.

42. Sarkar S, Barnaby R, Nymon AB, Taatjes DJ, Kelley TJ, Stanton BA. Extracellular vesicles secreted by primary human bronchial epithelial cells reduce Pseudomonas aeruginosa burden and inflammation in cystic fibrosis mouse lung. Am J Physiol Lung Cell Mol Physiol 326: L164–L174, 2024. doi: 10.1152/ajplung.00253.2023.

43. Gilles M-E, Slack FJ. Let-7 microRNA as a potential therapeutic target with implications for immunotherapy. Expert Opin Ther Targets 22: 929–939, 2018. doi: 10.1080/14728222.2018.1535594.

44. Schulte LN, Eulalio A, Mollenkopf H-J, Reinhardt R, Vogel J. Analysis of the host microRNA response to Salmonella uncovers the control of major cytokines by the let-7 family. EMBO J 30: 1977–1989, 2011. doi: 10.1038/emboj.2011.94.

45. Teng G, Wang W, Dai Y, Wang S, Chu Y, Li J. Let-7b is involved in the inflammation and immune responses associated with Helicobacter pylori infection by targeting Toll-like receptor 4. PLoS One 8: e56709, 2013. doi: 10.1371/journal.pone.0056709.

46. Reis M, Mavin E, Nicholson L, Green K, Dickinson AM, Wang X-N. Mesenchymal Stromal Cell-Derived Extracellular Vesicles Attenuate Dendritic Cell Maturation and Function. Front Immunol 9: 2538, 2018. doi: 10.3389/fimmu.2018.02538.

47. Home | ClinicalTrials.gov [Online]. [date unknown]. https://clinicaltrials.gov/ [22 Dec. 2024].

48. Sarkar S, Barnaby R, Nymon AB, Charpentier LA, Taub L, Wargo MJ, Weiss DJ, Bonfield TL, Stanton BA. Mesenchymal stromal cell extracellular vesicles reduce Pseudomonas biofilm formation, and let-7b-5p loading confers additional anti-inflammatory effects. Am J Physiol Lung Cell Mol Physiol 329: L455–L469, 2025. doi: 10.1152/ajplung.00187.2025.

49. IntaRNA: efficient prediction of bacterial sRNA targets incorporating target site accessibility and seed regions [Online]. [date unknown]. https://academic.oup.com/bioinformatics/article/24/24/2849/196716 [7 Feb. 2025].

50. Mann M, Wright PR, Backofen R. IntaRNA 2.0: enhanced and customizable prediction of RNA-RNA interactions. Nucleic Acids Res 45: W435–W439, 2017. doi: 10.1093/nar/gkx279.

51. Huang DW, Sherman BT, Lempicki RA. Systematic and integrative analysis of large gene lists using DAVID bioinformatics resources. Nat Protoc 4: 44–57, 2009. doi: 10.1038/nprot.2008.211.

52. Sherman BT, Hao M, Qiu J, Jiao X, Baseler MW, Lane HC, Imamichi T, Chang W. DAVID: a web server for functional enrichment analysis and functional annotation of gene lists (2021 update). Nucleic Acids Res 50: W216–W221, 2022. doi: 10.1093/nar/gkac194.

53. Zhu X, Badawi M, Pomeroy S, Sutaria DS, Xie Z, Baek A, Jiang J, Elgamal OA, Mo X, Perle KL, Chalmers J, Schmittgen TD, Phelps MA. Comprehensive toxicity and immunogenicity studies reveal minimal effects in mice following sustained dosing of extracellular vesicles derived from HEK293T cells. J Extracell Vesicles 6: 1324730, 2017. doi: 10.1080/20013078.2017.1324730.

54. Duan L, Huang H, Zhao X, Zhou M, Chen S, Wang C, Han Z, Han Z-C, Guo Z, Li Z, Cao X. Extracellular vesicles derived from human placental mesenchymal stem cells alleviate experimental colitis in mice by inhibiting inflammation and oxidative stress. International Journal of Molecular Medicine 46: 1551–1561, 2020. doi: 10.3892/ijmm.2020.4679.

55. Attaluri S, Jaimes Gonzalez J, Kirmani M, Vogel AD, Upadhya R, Kodali M, Madhu LN, Rao S, Shuai B, Babu RS, Huard C, Shetty AK. Intranasally administered extracellular vesicles from human induced pluripotent stem cell-derived neural stem cells quickly incorporate into neurons and microglia in 5xFAD mice. Front Aging Neurosci 15: 1200445, 2023. doi: 10.3389/fnagi.2023.1200445.

56. Rosenjack J, Hodges CA, Darrah RJ, Kelley TJ. HDAC6 depletion improves cystic fibrosis mouse airway responses to bacterial challenge. Sci Rep 9: 10282, 2019. doi: 10.1038/s41598-019-46555-4.

57. R-FAQ.pdf [Online]. [date unknown]. https://cran.r-project.org/doc/manuals/R-FAQ.pdf [22 Dec. 2024].

58. Koeppen K, Hampton TH, Barnaby R, Roche C, Gerber SA, Goo YA, Cho B-K, Vermilyea DM, Hogan DA, Stanton BA. An rRNA fragment in extracellular vesicles secreted by human airway epithelial cells increases the fluoroquinolone sensitivity of P. aeruginosa. Am J Physiol Lung Cell Mol Physiol 325: L54–L65, 2023. doi: 10.1152/ajplung.00150.2022.

59. Welsh JA, Goberdhan DCI, O’Driscoll L, Buzas EI, Blenkiron C, Bussolati B, Cai H, Di Vizio D, Driedonks TAP, Erdbrügger U, Falcon-Perez JM, Fu Q-L, Hill AF, Lenassi M, Lim SK, Mahoney MG, Mohanty S, Möller A, Nieuwland R, Ochiya T, Sahoo S, Torrecilhas AC, Zheng L, Zijlstra A, Abuelreich S, Bagabas R, Bergese P, Bridges EM, Brucale M, Burger D, Carney RP, Cocucci E, Crescitelli R, Hanser E, Harris AL, Haughey NJ, Hendrix A, Ivanov AR, Jovanovic-Talisman T, Kruh-Garcia NA, Ku’ulei-Lyn Faustino V, Kyburz D, Lässer C, Lennon KM, Lötvall J, Maddox AL, Martens-Uzunova ES, Mizenko RR, Newman LA, Ridolfi A, Rohde E, Rojalin T, Rowland A, Saftics A, Sandau US, Saugstad JA, Shekari F, Swift S, Ter-Ovanesyan D, Tosar JP, Useckaite Z, Valle F, Varga Z, van der Pol E, van Herwijnen MJC, Wauben MHM, Wehman AM, Williams S, Zendrini A, Zimmerman AJ, MISEV Consortium, Théry C, Witwer KW. Minimal information for studies of extracellular vesicles (MISEV2023): From basic to advanced approaches. J Extracell Vesicles 13: e12404, 2024. doi: 10.1002/jev2.12404.

60. Pegtel DM, Gould SJ. Exosomes. Annu Rev Biochem 88: 487–514, 2019. doi: 10.1146/annurev-biochem-013118-111902.

61. Ferguson SW, Wang J, Lee CJ, Liu M, Neelamegham S, Canty JM, Nguyen J. The microRNA regulatory landscape of MSC-derived exosomes: a systems view. Sci Rep 8: 1419, 2018. doi: 10.1038/s41598-018-19581-x.

62. Cavallaro S, Hååg P, Viktorsson K, Krozer A, Fogel K, Lewensohn R, Linnros J, Dev A. Comparison and optimization of nanoscale extracellular vesicle imaging by scanning electron microscopy for accurate size-based profiling and morphological analysis. Nanoscale Adv 3: 3053–3063, 2021. doi: 10.1039/d0na00948b.

63. Morente-López M, Fafián-Labora JA, Carrera M, de Toro FJ, Gil C, Mateos J, Arufe MC. Mesenchymal Stem Cell-Derived Extracellular Vesicle Isolation and Their Protein Cargo Characterization. Methods Mol Biol 2259: 3–12, 2021. doi: 10.1007/978-1-0716-1178-4_1.

64. Serrano-Pertierra E, Oliveira-Rodríguez M, Matos M, Gutiérrez G, Moyano A, Salvador M, Rivas M, Blanco-López MC. Extracellular Vesicles: Current Analytical Techniques for Detection and Quantification. Biomolecules 10: 824, 2020. doi: 10.3390/biom10060824.

65. Jensen OA, Prause JU, Laursen H. Shrinkage in preparatory steps for SEM. A study on rabbit corneal endothelium. Albrecht Von Graefes Arch Klin Exp Ophthalmol 215: 233–242, 1981. doi: 10.1007/BF00407662.

66. Saraiva M, O’Garra A. The regulation of IL-10 production by immune cells. Nat Rev Immunol 10: 170–181, 2010. doi: 10.1038/nri2711.

67. Barald KF, Shen Y-C, Bianchi LM. Chemokines and cytokines on the neuroimmunoaxis: Inner ear neurotrophic cytokines in development and disease. Prospects for repair? Exp Neurol 301: 92–99, 2018. doi: 10.1016/j.expneurol.2017.10.009.

68. Liu C, Chu D, Kalantar-Zadeh K, George J, Young HA, Liu G. Cytokines: From Clinical Significance to Quantification. Adv Sci (Weinh) 8: 2004433, 2021. doi: 10.1002/advs.202004433.

69. Caretti A, Peli V, Colombo M, Zulueta A. Lights and Shadows in the Use of Mesenchymal Stem Cells in Lung Inflammation, a Poorly Investigated Topic in Cystic Fibrosis. Cells 9: 20, 2019. doi: 10.3390/cells9010020.

70. Hampton TH, Thomas D, van der Gast C, O’Toole GA, Stanton BA. Mild Cystic Fibrosis Lung Disease Is Associated with Bacterial Community Stability. Microbiol Spectr 9: 10.1128/spectrum.00029-21, [date unknown]. doi: 10.1128/spectrum.00029-21.

71. Litman PM, Day A, Kelley TJ, Darrah RJ. Serum inflammatory profiles in cystic fibrosis mice with and without Bordetella pseudohinzii infection. Sci Rep 11: 17535, 2021. doi: 10.1038/s41598-021-97033-9.

72. Varderidou-Minasian S, Lorenowicz MJ. Mesenchymal stromal/stem cell-derived extracellular vesicles in tissue repair: challenges and opportunities. Theranostics 10: 5979–5997, 2020. doi: 10.7150/thno.40122.

73. Pang Z, Raudonis R, Glick BR, Lin T-J, Cheng Z. Antibiotic resistance in Pseudomonas aeruginosa: mechanisms and alternative therapeutic strategies. Biotechnol Adv 37: 177–192, 2019. doi: 10.1016/j.biotechadv.2018.11.013.

74. Cruz FF, Rocco PRM. The potential of mesenchymal stem cell therapy for chronic lung disease. Expert Rev Respir Med 14: 31–39, 2020. doi: 10.1080/17476348.2020.1679628.

75. Tognon M, Köhler T, Luscher A, van Delden C. Transcriptional profiling of Pseudomonas aeruginosa and Staphylococcus aureus during in vitro co-culture. BMC Genomics 20: 30, 2019. doi: 10.1186/s12864-018-5398-y.

76. Cendra MDM, Torrents E. Pseudomonas aeruginosa biofilms and their partners in crime. Biotechnol Adv 49: 107734, 2021. doi: 10.1016/j.biotechadv.2021.107734.

77. Van Deun J, Mestdagh P, Agostinis P, Akay Ö, Anand S, Anckaert J, Martinez ZA, Baetens T, Beghein E, Bertier L, Berx G, Boere J, Boukouris S, Bremer M, Buschmann D, Byrd JB, Casert C, Cheng L, Cmoch A, Daveloose D, De Smedt E, Demirsoy S, Depoorter V, Dhondt B, Driedonks TAP, Dudek A, Elsharawy A, Floris I, Foers AD, Gärtner K, Garg AD, Geeurickx E, Gettemans J, Ghazavi F, Giebel B, Kormelink TG, Hancock G, Helsmoortel H, Hill AF, Hyenne V, Kalra H, Kim D, Kowal J, Kraemer S, Leidinger P, Leonelli C, Liang Y, Lippens L, Liu S, Lo Cicero A, Martin S, Mathivanan S, Mathiyalagan P, Matusek T, Milani G, Monguió-Tortajada M, Mus LM, Muth DC, Németh A, Nolte-’t Hoen ENM, O’Driscoll L, Palmulli R, Pfaffl MW, Primdal-Bengtson B, Romano E, Rousseau Q, Sahoo S, Sampaio N, Samuel M, Scicluna B, Soen B, Steels A, Swinnen JV, Takatalo M, Thaminy S, Théry C, Tulkens J, Van Audenhove I, van der Grein S, Van Goethem A, van Herwijnen MJ, Van Niel G, Van Roy N, Van Vliet AR, Vandamme N, Vanhauwaert S, Vergauwen G, Verweij F, Wallaert A, Wauben M, Witwer KW, Zonneveld MI, De Wever O, Vandesompele J, Hendrix A. EV-TRACK: transparent reporting and centralizing knowledge in extracellular vesicle research. Nat Methods 14: 228–232, 2017. doi: 10.1038/nmeth.4185.

78. Perez-Riverol Y, Bandla C, Kundu DJ, Kamatchinathan S, Bai J, Hewapathirana S, John NS, Prakash A, Walzer M, Wang S, Vizcaíno JA. The PRIDE database at 20 years: 2025 update. Nucleic Acids Res 53: D543–D553, 2025. doi: 10.1093/nar/gkae1011.

